# LEGEND: Identifying Co-expressed Genes in Multimodal Transcriptomic Sequencing Data

**DOI:** 10.1101/2024.10.27.620451

**Authors:** Tao Deng, Mengqian Huang, Kaichen Xu, Yan Lu, Yucheng Xu, Siyu Chen, Nina Xie, Hao Wu, Xiaobo Sun

**Author notes:** Corresponding authors. (Sun X), (Wu H). Equal contribution (Tao Deng, Xiaobo Sun).

## Abstract

Identifying co-expressed genes across tissue domains and cell types is essential for revealing co-functional genes involved in biological or pathological processes. While both single-cell RNA-sequencing (scRNA-seq) and spatially-resolved transcriptomic (SRT) data offer insights into gene co-expression patterns, current methods typically utilize either data type alone, potentially diluting the co-functionality signals within co-expressed gene groups. To bridge this gap, we introduce LEGEND, a novel computational method that integrates scRNA-seq and SRT data for identifying groups of co-expressed genes at both cell type and tissue domain levels. LEGEND employs an innovative hierarchical clustering algorithm designed to maximize intra-cluster redundancy and inter-cluster complementarity, effectively capturing more nuanced patterns of gene co-expression and spatial coherence. Enrichment and cofunction analyses further showcase the biological relevance of these gene clusters, and their utilities in exploring context-specific novel gene functions. Notably, LEGEND can reveal shifts in gene-gene interactions under different conditions, furnishing insights for disease-associated gene crosstalk. Moreover, LEGEND can be utilized to enhance the annotation accuracy of both spatial spots in SRT and single-cells in scRNA-seq, and pioneers in identifying genes with designated spatial expression patterns. LEGEND is available at https://github.com/ToryDeng/LEGEND.

## Introduction

Single-cell RNA sequencing (scRNA-seq) technology allows the profiling of whole-transcriptome at individual cell resolution, yielding rich information on transcriptional activities among diverse cell types and gene expression dynamics in complex tissues and diseases [1–3]. However, the scRNA-seq data lacks spatial information of single-cells, which hinders the understanding of gene functions in the context of tissue microenvironment. Spatially-resolved transcriptomics (SRT) addresses this issue by profiling spatial gene expression in tissues, thus providing unprecedented opportunities to characterize spatial distribution of cell types [4], delineate spatial tissue organization [5], gene-gene interactions [6], etc. Nonetheless, current high-throughput SRT technologies typically have to trade single-cell resolution for full transcriptome coverage (e.g., in situ capturing-based technologies such as 10x Visium [7]), making it difficult to reliably determine single cell identities at each measurement site (i.e., spatial spot). Therefore, the integrative analysis of SRT and scRNA-seq data that utilizes their complementary information can significantly facilitate analytical tasks such as identifying gene expression patterns at both tissue domain and cell type levels.

The identification of gene expression patterns is essential for revealing gene functions and interactions involved in biological processes including cell differentiation and tissue development. In the context of scRNA-seq and SRT, researchers focus on three categories of gene expression patterns: differentially expressed genes (DEGs), variably expressed genes including highly variable genes (HVGs) in scRNA-seq and spatially variable genes (SVGs) in SRT, and co-expressed genes [5,8]. They provide different insights into the underlying biological processes and genetic mechanisms. For instance, DEGs facilitate the discovery of cell type-/tissue region-specific biomarkers: Variably expressed genes allow the detection of genes involved in the differentiation of cell types and tissue domains [8,9]: Co-expressed genes can reveal functionally related genes and pathways [10,11], providing genetic clues to disease pathogenetic mechanisms based on their co-expression patterns associated with the development of pathology [12]. Intriguingly, it has been suggested co-expressed genes can serve as genomic “contexts” for learning distributed gene representations to facilitate downstream analytical tasks, similar to the word context used in word2vec [13,14].

A variety of computational methods have been developed for identifying gene expression patterns in scRNA-seq and SRT [9,15–18]. Specifically, methods for identifying co-expressed genes include scGeneClust [11], CS-CORE [19], and COTAN [20] for scRNA-seq, and CNN-Preg [10], Giotto [15], STUtility [21], and SPARK [22] for SRT. However, these methods are constrained by several limitations. First, they work on SRT or scRNA-seq data alone and analyzing gene co-expression over cell types or tissue domains alone. This leads to the identification of weaker co-functional genes in that they may exhibit similar expression patterns across tissue domains but not across cell types or vice versa. Second, all methods for SRT except CNN-PReg evaluate gene expression similarity for each individual spot independently, neglecting the spatial relationship among spatial spots and the overall spatial patterns. Third, these methods fall short of offering means to leverage identified co-expressed genes for downstream applications, such as pinpointing genes with targeted spatial expression patterns or improving the information efficiency of feature genes used by analytical algorithms.

In response, we develop a novel method, mu**L**timodal co-**E**xpressed **GEN**es fin**D**er (LEGEND), which identifies co-expressed genes across both cell types and tissue domains under the framework of information theory. LEGEND estimates gene relevance, redundancy, and complementarity in both SRT and scRNA-seq datasets in a pseudo-semi-supervised manner. This information is used to construct a gene-gene redundancy graph, upon which genes are grouped as per their redundancy into hierarchical clusters, which represent co-expressed and cofunctional gene modules. We evaluate LEGEND over SRT and scRNA-seq datasets from adult mouse brain and human dorsolateral prefrontal cortex (DLPFC) (**Table 1**) and find that genes within the same cluster exhibit similar spatial expression patterns across all SRT datasets. In comparison to seven existing methods (**Table 2**) and an adapted variant of LEGEND (referred as LEGEND-no-sc) that omits scRNA-seq data usage, LEGEND shows superior gene clustering performance in terms of gene co-expression across cell types and spatial coherence across tissue domains. Additionally, via pathway enrichment analysis and gene cofunction analysis, we find that LEGEND effectively groups context-specific cofunctional genes into clusters, with aggregated expression patterns overlapping with tissue anatomical architectures and pathological distributions. Furthermore, we demonstrate LEGEND’s utilization in three key downstream tasks including identifying disease-associated differential gene co-expressions, pinpointing genes with designated spatial expression patterns, and improving the accuracy of both cell clustering in scRNA-seq and spatial clustering in SRT. Real data analyses showcase our method’s superior performance across all these tasks and promise its contributions in revealing gene-level pathogenic mechanisms.

**Table 1.**
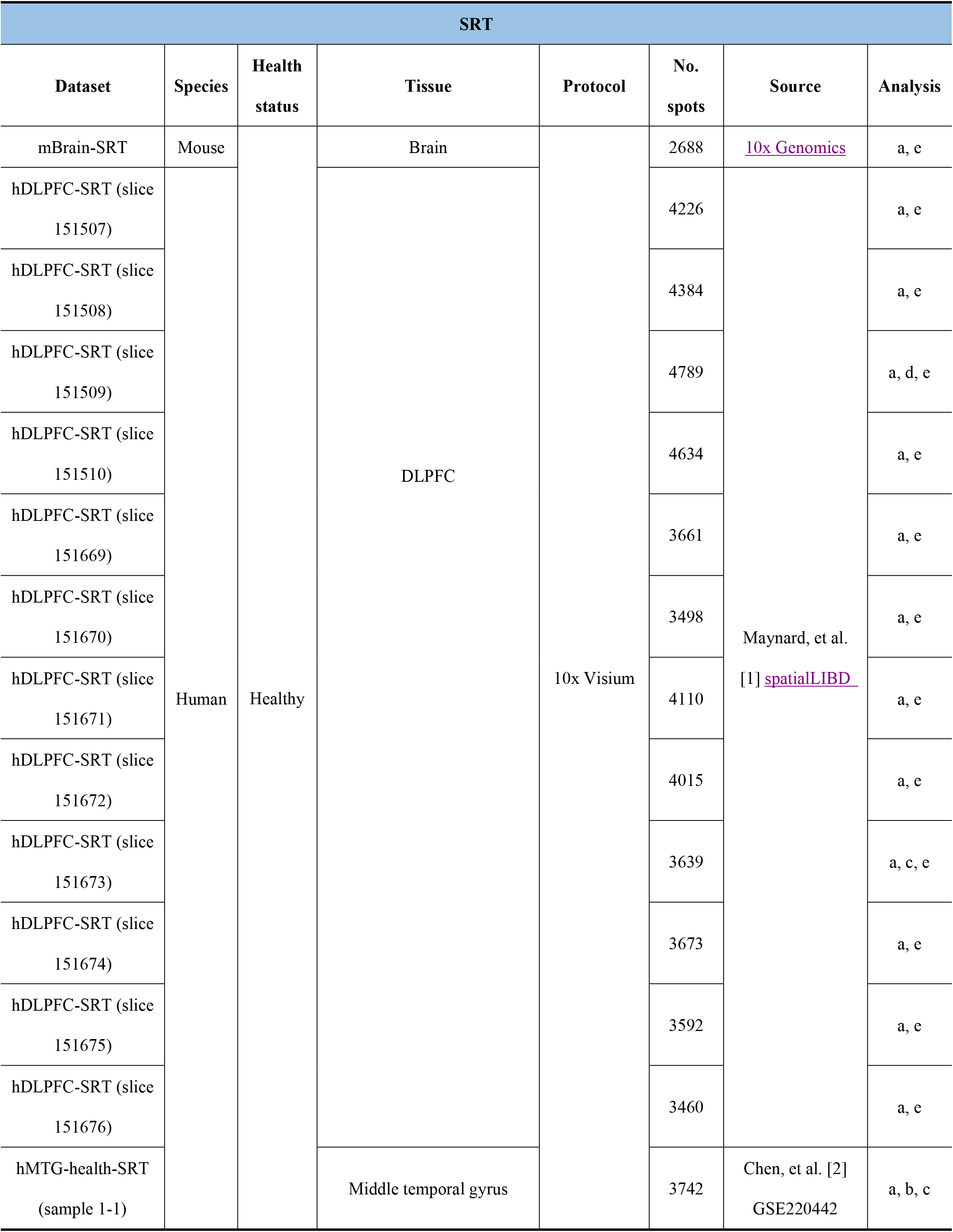
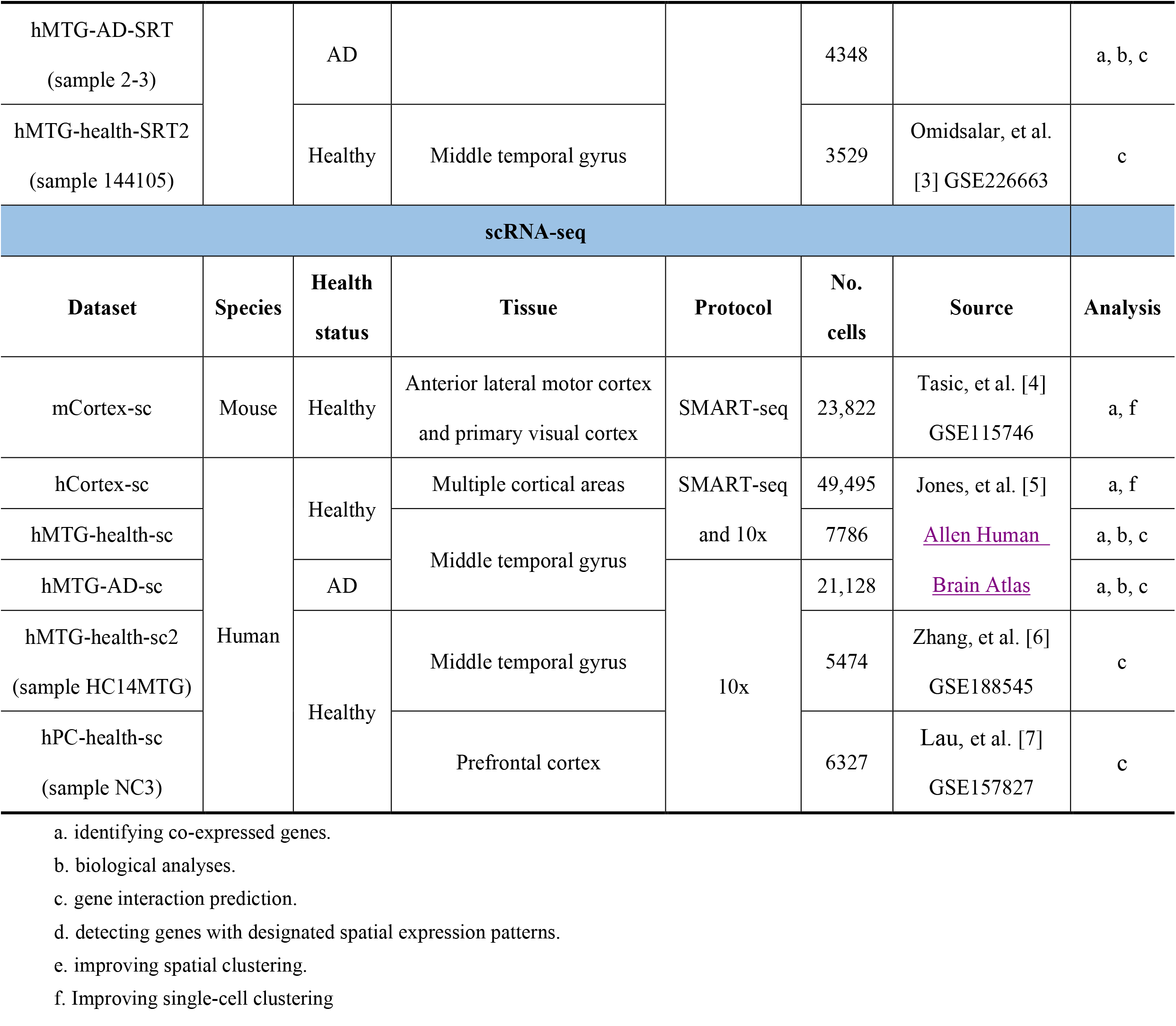
Datasets used in this study.

**Table 2.** List of benchmark methods.

| Method | Target data type | Core methodology | Implementation | Ref. | Analysis |
| --- | --- | --- | --- | --- | --- |
| scGeneClust | scRNA-seq | Splitting information-based maximum-spanning tree | Python package *GeneClust* 0.0.1 | [1] | Identifying co-expressed genes |
| CS-CORE |  | Hierarchical clustering | Python package *CSCORE* 1.0.0 | [2] |  |
| COTAN |  | Hierarchical clustering | R package *COTAN* 2.5.0 | [3] |  |
| CNN-PReg | SRT | CNN with graph regularization | https://github.com/kuanglab/CNN-PReg, commit 5a8505d (no available package) | [4] |  |
| Giotto |  | Hierarchical clustering | R package *Giotto* 3.2 | [5] |  |
| STUtility |  | Non-negative matrix factorization | R package *STUtility* 1.1.1 | [6] |  |
| SPARK |  | Hierarchical clustering | R package *SPARK* 1.1.1 | [7] |  |
| CS-CORE | scRNA-seq | The joint distribution over UMI counts | Python package *CSCORE* 1.0.0 | [2] | Predicting gene interactions |
| COTAN |  | A statistic based on contingency tables of binarized UMI counts | R package *COTAN* 2.5.0 | [3] |  |
| Giotto | SRT | Pearson correlation | R package *Giotto* 3.2 | [5] |  |
| scGeneClust | scRNA-seq | Feature relevance | Python package *GeneClust* 0.0.1 | [1] | Selecting feature genes |
| VST |  | Variances of standardized counts | Python package *scanpy* 1.9.3 | [8] |  |
| M3Drop |  | Differences between gene-specific parameters and the global parameter | R package *M3Drop* 1.12.0 | [9] |  |
| SpatialDE | SRT | Likelihood ratio tests based on Gaussian process regression | Python package *SpatialDE* 1.1.3 | [10] |  |
| SPARK-X |  | Non-parametric covariance tests | R package *SPARK* 1.1.1 | [7] |  |
| BinSpect (kmeans) |  | Fisher tests on gene expression levels binarized with k-means | R package *Giotto* 3.2 | [5] |  |
| BinSpect (rank) |  | Fisher tests on gene expression levels binarized with threshold ranking |  |  |  |

## Materials and methods

### Data preprocessing

For both scRNA-seq and SRT data, we adhere to the standard pipeline of data preprocessing provided by the SCANPY package [23] which involves several steps: Firstly, we filter out mitochondrial and External RNA Controls Consortium (ERCC) spike-in genes. Secondly, genes detected in fewer than 10 cells in scRNA-seq datasets or 10 spots in SRT datasets are excluded. Cells with fewer than 200 detected genes in scRNA-seq datasets are also removed. We do not perform filtering on spatial spots in SRT datasets to maintain spatial data integrity. Lastly, the gene expression counts of both scRNA-seq and SRT datasets are normalized by library size, followed by log-transformation.

### The algorithm of LEGEND

LEGEND’s framework is structured into two primary stages that focus on cell type and tissue domain pseudo-labelling, and on gene clustering, as illustrated in **Figure 1** and **Table 3**. The core idea of LEGEND is that co-expressed and co-functional genes typically exhibit similar expression patterns in both scRNA-seq and SRT datasets, which can be identified through integrated gene clustering. In Stage I, LEGEND begins by generating highly confident pseudo-labels of spatial domains for SRT data, as detailed in the following section. The pseudo-labelling of single-cells in scRNA-seq is discussed in **Supplementary Text S1**. In Stage II, these pseudo-labels are utilized to estimate the similarity and discriminability of gene expression patterns across cell types and tissue domains in terms of relevance, redundancy, and complementarity from information theory (see “*Gene relevance, redundancy and complementarity*” section below). Next, gene relationships are embedded in a gene redundancy graph, where nodes represent individual genes and edge weights indicate the redundancy between connected gene pairs. Leveraging a novel hierarchical clustering algorithm, LEGEND assigns informationally redundant genes to the same groups, while separating informationally complementary ones into distinct groups so as to maximize intra-group redundancy and inter-group complementarity. As such, these groups represent functionally distinct gene modules, each comprising genes that are co-expressed and cofunctional across cell types and tissue domains.

**Table 3.**
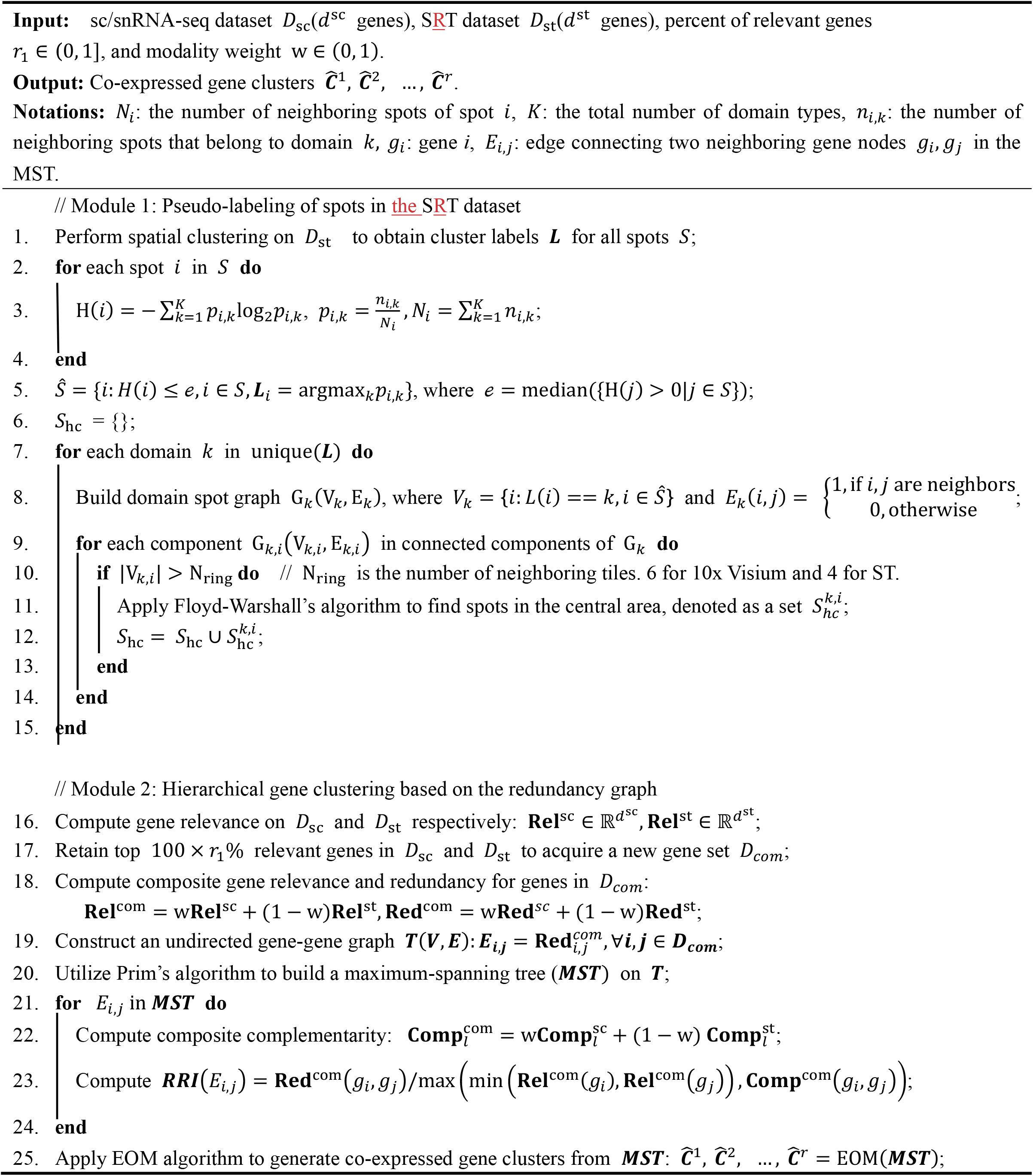
Algorithm of LEGEND.

**Figure 1.**
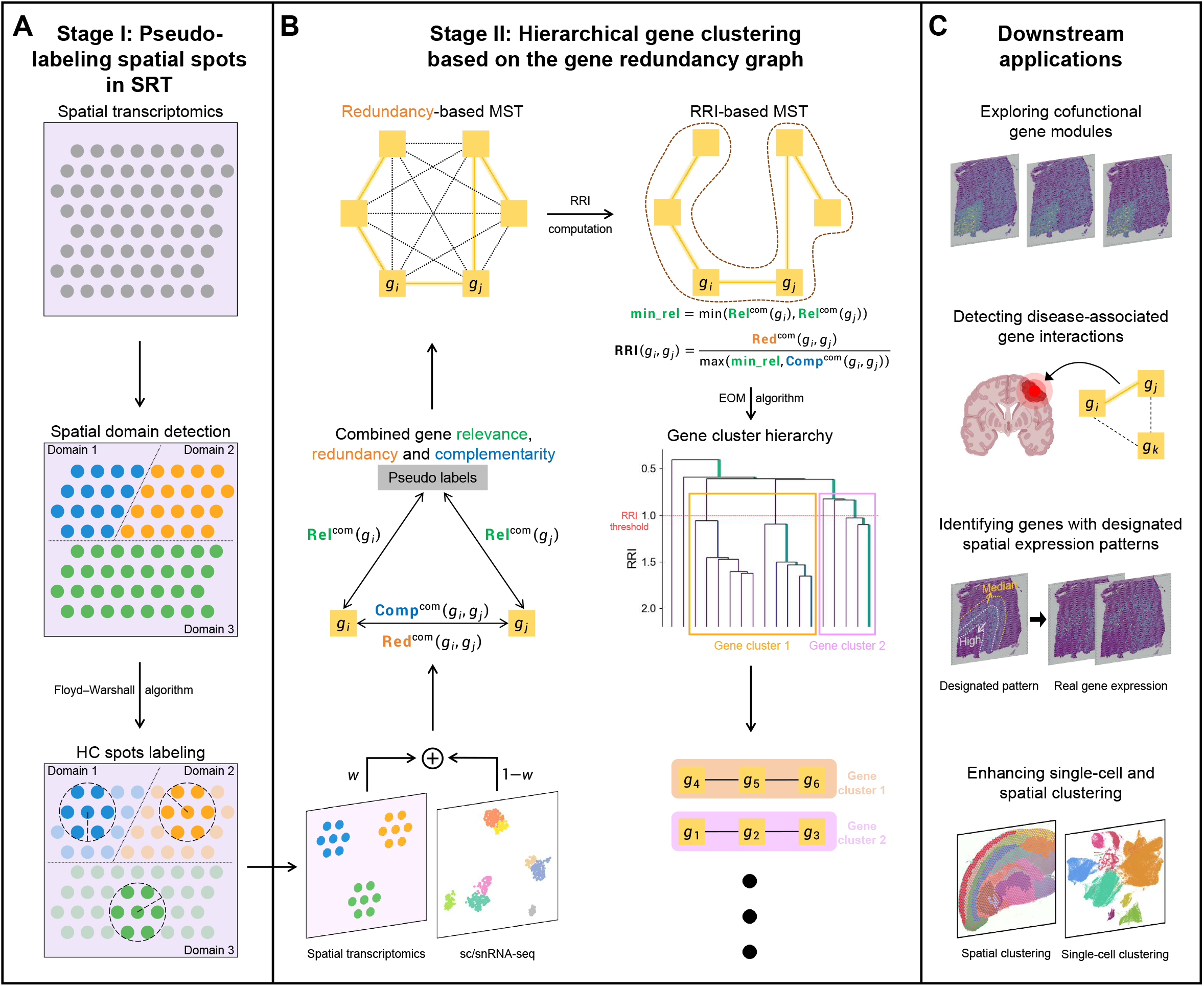
The workflow of LEGEND. It consists of two stages: **A**. Pseudo-labelling spatial spots in SRT. LEGEND initially groups spatial spots into spatial domains using an off-the-shelf spatial clustering algorithm. Floyd-Warshall’s algorithm is then employed to identify domain central areas wherein spots (i.e., highly confident (HC) spots) reliably belong to the same domain. HC spots are used in subsequent steps while domain marginal spots are excluded. **B**. Hierarchical gene clustering based on the gene redundancy graph. Gene relevance, redundancy, and complementarity are computed in both SRT and scRNA-seq datasets based on pseudo-labels of HC spots and HC cells (if true single-cell labels are unavailable). These values are then combined as weighted average composite values. Then, a complete gene redundancy graph is constructed, with edge weights equal to composite redundancy between connecting gene pairs. From this graph, an MST is extracted and converted into a hierarchy of gene clusters through the EOM algorithm. Within this hierarchy, high-quality gene clusters, representing groups of co-expressed genes, are automatically determined based on their cluster stability coefficients. **C**. Gene clustering-based downstream applications. LEGEND-mediated gene clustering allows exploring cofunctional gene modules, detecting disease-associated gene interactions, pinpointing genes with designated spatial expression patterns, and enhancing both single-cell and spatial clustering.

### Pseudo-labelling of spatial spots in SRT

In this stage, a pseudo-semi-supervised approach is adopted to acquire pseudo-labels for spatial spots that are likely to be of the same domain types, termed highly confident (HC) spots. Stage I unfolds in two steps (Figure 1A). Initially, spatial spots are clustered into spatial domains using an off-the-shelf spatial clustering method, such as SpaGCN, from which HC spots are selected. Since there is greater confidence in the cluster labels of spots at the center of large, homogeneous spatial domains, peripheral spots are excluded upfront. A spot is deemed peripheral if its neighboring spot labels are significantly different from the spot’s label or show considerable heterogeneity. Therefore, a spot *i* is excluded if the labels of its neighboring spots are predominantly different or if the label distribution exhibits high entropy, which is defined as:

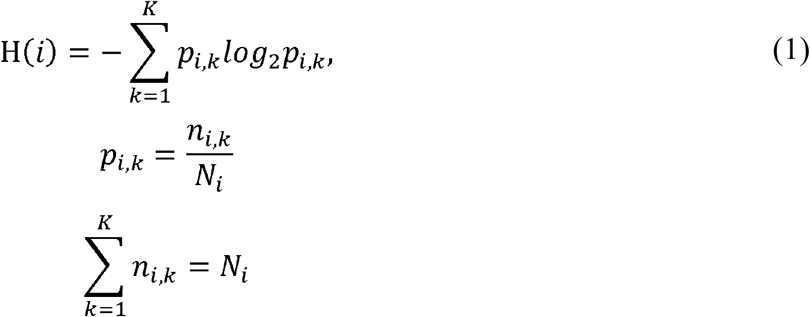

where *N*_*i*_ is the number of neighboring spots of spot *i*, is the total number of domain types, *n*_*i*,*k*_ is the number of neighboring spots that belong to domain type *k* ∈{1,…, *K*}, and p_*i*,*k*_ is the proportion of neighboring spots that belong to domain type *k*. Subsequently, we construct a graph *G*_*k*_(*V*_*k*_,*E*_*k*_) for each domain type *k*:

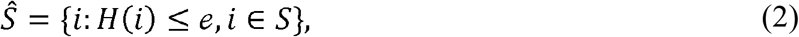

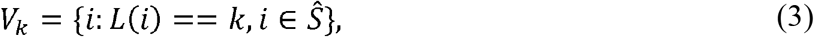

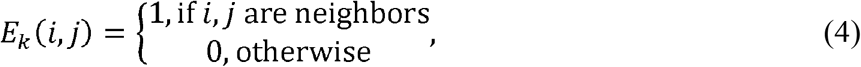

where *S* denotes the set of all spots, *Ŝ* the set of spots after entropy filtering, *L(i)* the domain type label of spot *i*. Note that, spots of the same domain type may locate in separated spatial regions, so the graph of a domain type may consist of multiple disconnected subgraphs. We exclude subgraphs whose sizes are smaller than the number of neighboring tiles (e.g., 6 for 10x Visium data). To identify HC spots, we first need to pinpoint the graph center for each remaining subgraph. Specifically, for a given spot u within a subgraph *G*_*k*,*i*_(*V*_*k*_,*E*_*k*_) ∈ G_*k*_(*V*_*k*_,*E*_*k*_), its eccentricity is computed as, 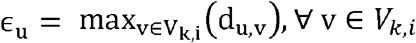. Here d_u,v_ denotes the distance between spots u and *v*, which is calculated using the Floyd-Warshall’s algorithm [24]. The center ψ(*G*_*k*,*i*_) and radius r(*G*_*k*,*i*_) of *G*_*k*,*i*_(*V*_*k*_,*E*_*k*_) are:

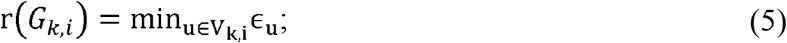

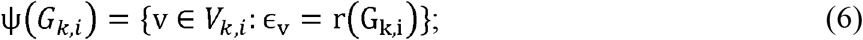

Spots whose distance to ψ(*G*_*k*,*i*_) is less than 0.6x r(*G*_*k*,*i*_) _*i*_ are deemed as close to the graph center and selected as HC spots of *G*_*k*,*i*_.

### Gene relevance, redundancy, and complementarity

Gene relevance, redundancy, and complementarity are computed as mutual information (MI) between gene-spot/cell label or gene-gene pairs. Given MI is originally defined for discrete variables and gene expression levels are continuous, we leverage a k-nearest neighbor-based algorithm (see Supplementary Text S2) when MI calculation involves continuous variables.

*Gene relevance*. The relevance of a gene reflects its discriminative power on different domain/cell type labels. Let **g**_i_ ∈ ℝ^n^ denote gene *i*’s expression level across spatial spots or cells, H(**g**_i_) its entropy, and H(**g**_i_|***L***) its conditional entropy given the labels. Gene *i*’s relevance is computed as:

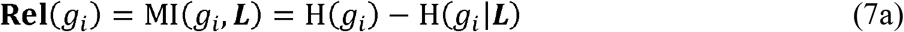

We further define the conditional relevance of gene *i* given *g*_*j*_ as:

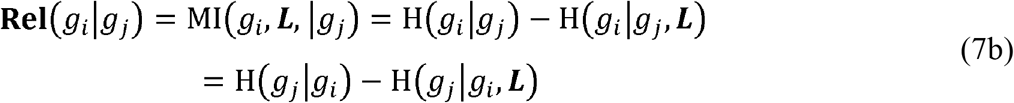

where H(*g*_i_|*g*_*j*_) and H(*g*_i_|*g*_*j*_,***L***) are conditional entropies.

#### Gene redundancy

The redundancy between a pair of genes *i* and 1 is defined as the MI between their expression levels:

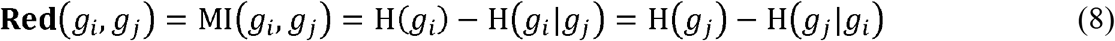

Gene redundancy represents the amount of information shared between genes, implying that discarding highly redundant genes has minimal impact on the overall information and enhances the informational efficiency of feature gene set.

#### Gene complementarity

The complementarity between genes *i* and *j* measures the additional information gained for label discrimination when both genes are considered together compared to each on its own:

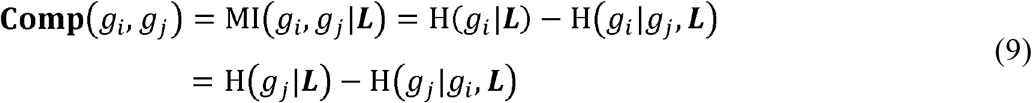

Furthermore, we have the following theorem (see Supplementary Text S3 for proof):

##### Theorem 1.

Red(*g*_i_|*g*_*j*_) < **Comp**(*g*_i_|*g*_*j*_) ⇔ **Rel**(*g*_i_) < **Rel**(*g*_i_|*g*_*j*_) & **Rel**(*g*_*j*_) < **Rel**(*g*_i_|*g*_*j*_)

It shows that if the complementarity of two genes is larger than their redundancy, the relevance of one gene can be enhanced given the expression levels of the other, i.e., **Rel**(*g*_i_|*g*_*j*_) > **Rel**(*g*_*j*_)

### Redundancy graph-based hierarchical gene clustering

Gene relevance, redundancy, and complementarity in the SRT and scRNA-seq datasets are computed using the labels of HC spots and single-cells. Only the most informative genes (i.e., the top 80% in relevance in both datasets) are retained. For two given genes *g*_*i*_ and *g*_*j*_, their composite gene relevance, redundancy, and complementarity (**Rel**^com^, **Rel**^com^ and **Rel**^com^) are computed as a weighted average of the individual metrics derived from the SRT and scRNA-seq datasets:

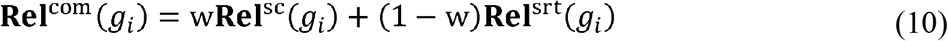

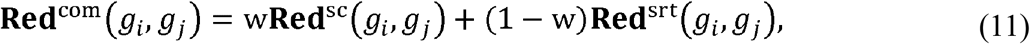

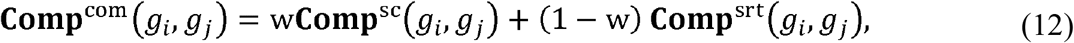

where w ∈ (0,1) is the weight factor for balancing the contribution of SRT and scRNA-seq data. Following this, an undirected complete gene graph is constructed, wherein vertices represent genes and edge weights equal the composite redundancies between connected gene pairs. The graph is partitioned into subgraphs representing gene clusters, aiming to maximize intra-cluster gene redundancy and inter-cluster complementarity. Instead of partitioning the complete graph, an NP-hard problem, we conduct the partition on a maximum-spanning tree (MST) extracted from the graph using Prim’s algorithm [25]. This MST not only significantly reduces the number of edges to be partitioned but also preserves the graph’s most representative redundancy structure. Formally, we first define the relative redundancy index (RRI) of the edge *e*_*i*,*j*_ connecting two neighboring genes *g*_*i*_,*g*_*j*_ on the MST as:

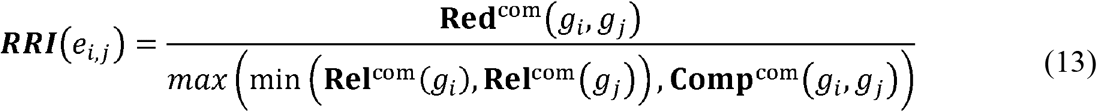

The EOM algorithm [26] is employed to partition the MST at different RRI thresholds: *e*_*i*,*j*_, is preserved only if ***RRI***(*e*_*i*,*j*_), is above the current RRI threshold, generating a hierarchy of gene clusters.

As the RRI threshold increases along the hierarchy in the top-down direction, fewer edges are preserved, leading to the formation of finer gene clusters toward the hierarchy’s bottom. Among all clusters within the hierarchy, clusters of high quality and sensible size are automatically determined based on their stability coefficients, which essentially measure their “persistence” along the hierarchy (see Supplementary Text S4). The maximum RRI threshold is set to 1 so that genes *i* and *j* are assigned to different clusters only if either of the following two inequalities is satisfied:

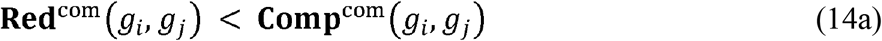

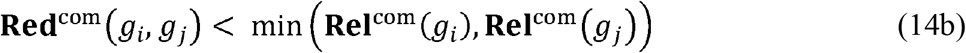

Inequality 14a implies that two genes with greater complementarity than their redundancy should be separated into different clusters to enhance their overall relevance, as suggested by Theorem 1 above.

Inequality 14b implies that if two genes should not be in the same cluster if they are not sufficiently redundant relative to their minimum relevance. These two conditions ensure the informational complementarity between different clusters. Furthermore, clusters with a size smaller than 5 are considered as noise groups of genes “falling out of their parent cluster” rather than “real clusters”, and merged back into their parent clusters in the hierarchy.

### Biological analysis

We investigate the biological significance of LEGEND-generated gene clusters through a case study involving two pairs of SRT and scRNA-seq datasets from the middle temporal gyrus (MTG) of both AD patients and healthy individuals. The analysis proceeds with the following steps:

#### Selection of top relevant gene clusters

Gene clusters are sorted in descending order by the maximum **Rel**^com^ of their component genes, from which top-ranked clusters are selected until they collectively encompass 100 genes. Gene clusters selected from the AD patient datasets constitute the disease group, while those from the healthy individual datasets form the normal group. Both groups are subjected to enrichment and co-function analysis. Concurrently, an equal number of genes are randomly selected as a control group.

#### Enrichment analysis

Biological processes enriched in the given gene set can be identified via enrichment analysis. We conduct both gene ontology (GO) and KEGG pathway enrichment analyses for disease, normal and control gene groups to assess their biological relevance to MTG functions and AD pathology. The R package clusterProfiler (v4.2.2) [27] is utilized to identify GOBPs or KEGG pathways enriched in the genes of the three groups. Enrichment resultant p-values are adjusted via the Benjamini-Hochberg procedure, and the 20 most significantly enriched GOBPs/KEGG pathways are selected for further biological analyses, as described in the Results section.

#### Gene co-function analysis

Functional coherence of a gene set within a GOBP or KEGG pathway can be assessed using the neighbor-voting method proposed by Ballouz et al [28]. The rationale is that if some genes’ involvements in GOBPs/pathways can be more accurately predicted by other genes’, then these genes exhibit greater functional coherence. Formally, let d denote the number of genes, q denote the number of GOBPs/pathways, Σ ∈ ℝ^*d*×*d*^ denote the gene-gene similarity matrix of the SRT dataset, 𝒮 ∈ ℝ^*d*×*d*^ denote the gene-gene similarity matrix of the scRNA-seq dataset, and **A** ∈ ℝ^*d*×*q*^ denote the gene annotation matrix. Then we have:

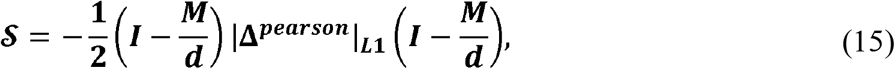

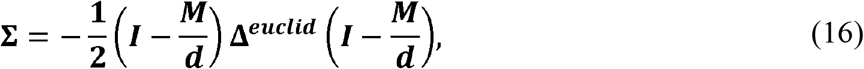

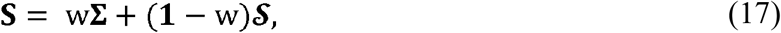

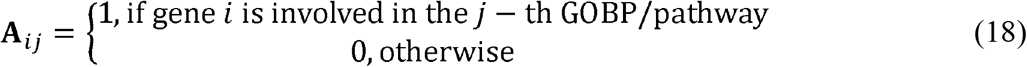

where *M* ∈ ℝ^*d*×*d*^ is a matrix of 1s, and **Δ**^*pearson*^, **Δ**^*euclid*^∈ ℝ^*d*×*d*^ are gene-gene distance matrices for the scRNA-seq and SRT datasets, as calculated from Equation 23a and 23b, respectively (see “*evaluation metrics*” section). S ∈ ℝ^*d*×*d*^ is a weighted average of Σ and 𝒮, with weight w assigned to Σ.

The prediction accuracy is assessed via three-fold cross validation conducted on the top 20 enriched GOBPs/pathways. In each fold, one third of the genes are randomly selected as the test dataset, and their associated GOBPs/pathways are masked by setting the corresponding rows in **A** to zeros. Given a gene, in the test dataset, its predicted probability of being involved in the *j*-th GOBP/pathway is calculated as:

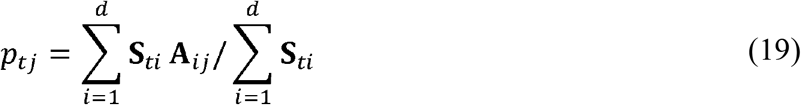

The prediction results are evaluated using AUC metric, where a higher value indicates greater functional coherence among the genes in that GOBP/pathway. Genes involved in the selected GOBPs and KEGG pathways are obtained using R package AnnotationDbi (v1.56.2) and KEGGREST (v1.34.0), respectively. The R package EGAD (v1.22.0) is employed to conduct the neighbor-voting-based prediction.

### Identifying disease-associated gene crosstalk

Given an scRNA-seq dataset 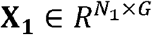 and an SRT dataset 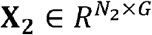, where *N*_l_ denotes the number of single-cells, *N*_2_ the number of spots, and *G* the number of genes, we first calculate its combined Pearson correlation matrix of genes Θ ∈ R^*G*×*G*^ = *w* × *corr*(**X**_1_) + (1−*w*) × *corr*(**X**_2_). Let Σ ∈ R^*G*×*G*^denote the gene-gene redundancy matrix generated by LEGEND. is used as a surrogate non-parametric covariance matrix in a Gaussian Copula Graphical Model (GCGM) to estimate the sparsified gene-gene precision matrix 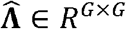:

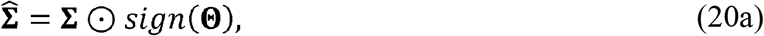

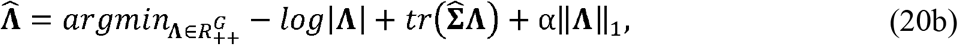

where ⊙ is the Hadamard Product. 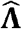 is then converted into the gene partial correlation matrix Φ ∈ R^*G*×*G*^, wherein nonzero values indicate plausible gene-gene interactions:

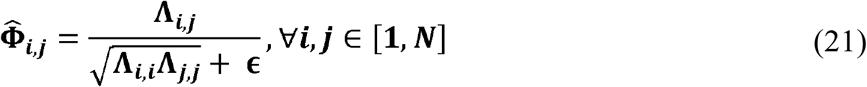

where ∈ = 1*e* − 6 is for numerical stability. In our study, we compute two gene partial correlation matrices, Φ_*AD*_ and Φ_*health*_, which are derived from AD and healthy datasets, respectively. The absolute partial correlation difference matrix Φ_*diff*_: = |Φ_*AD*_ − Φ_*health*_| can reveal disease-associated gene-gene interactions (i.e., gene pairs corresponding to largest absolute partial correlation shifts).

### Evaluation metrics

#### Gene clustering

Davies-Bouldin (DB) index [29] is used to assess the performance of gene clustering in SRT and scRNA-seq:

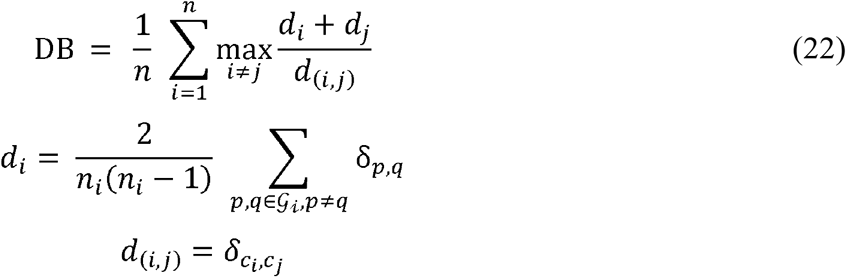

where δ_*p*,*q*_, denotes the distance between genes *p* and *q, n* denotes the number of clusters, d_*i*_ is the intra-cluster distance of cluster *i* (calculated as the average distance between genes of cluster *i*), and *d*_(*i*,*j*)_ is the inter-cluster distance between clusters *i* and *j* (calculated as the distance between cluster centroids *c*_*i*_ and *c*_*j*_). DB index represents the ratio of intra-cluster distance (compactness) to inter-cluster distance (separateness), so a smaller value of DB index indicates better clustering performance. To evaluate the co-expression of gene clusters in the scRNA-seq dataset and the spatial coherence of gene clusters in the SRT dataset, Pearson distance and Euclidean distance are used in the calculation of DB index, respectively [10]:

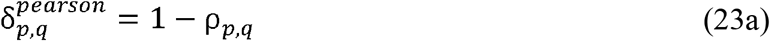

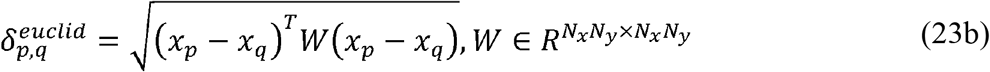

where ρ_*p*,*q*_, represents the Pearson correlation between genes *p* and *q* in the scRNA-seq dataset. *N*_*x*_and *N*_*y*_ denote the number of spatial spots along the horizontal and vertical axes of the spatial map of the SRT dataset, respectively. *x*_*p*_ is the flattened expression matrix of gene *p* in the SRT dataset, and *W* is the weight matrix calculated based on the spatial spots’ locations using a Gaussian kernel, reflecting the spatial closeness of the spatial spots.

#### Spatial spots/single-cell clustering

The accuracy of clustering for spatial spots/cells is assessed using the Adjusted Rand Index (ARI) and Normalized Mutual Information (NMI). Let denote the total number of spots/cells, *n*_*ij*_ the number of spots/cells of domain/cell type *i* in cluster *j*, *a*_*i*_ the total number of spots/cells of domain/cell type *i*, *b*_*j*_ the number of all spots/cells in cluster *j*. Additionally, *L* refers to the true labels of spots/cells, and 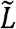 refers to the csluster labels of spots/cells. The ARI and NMI are defined as follows:

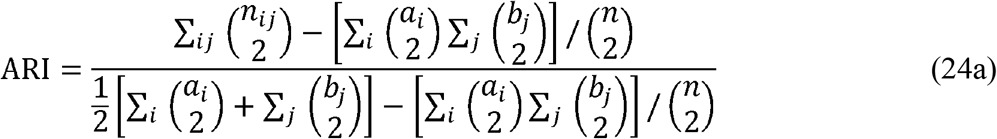

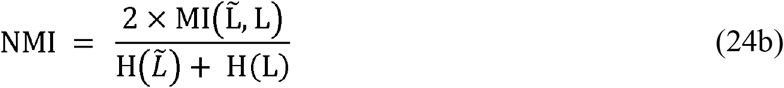

where *MI* represents mutual information and H represents entropy.

## RESULTS

### LEGEND effectively identifies cross-modal clusters of co-expressed genes

In this section, we compare the clustering performance of LEGEND against six state-of-the-art (SOTA) competing methods (CNN-PReg, Giotto, CS-CORE, COTAN, SPARK, and STUtility) in identifying co-expressed gene clusters. We also include scGeneClust and LEGEND-no-sc, both of which uses scRNA-seq data only, as competing methods. Our analysis involves 12 SRT datasets from the human DLPFC (hDLPFC-SRT), all paired with a scRNA-seq dataset (hCortex-sc) from multiple human cortical areas: one SRT dataset from the adult mouse brain (mBrain-SRT) accompanied by a corresponding scRNA-seq dataset (mCortex-sc): one SRT dataset (hMTG-health-SRT) and a matching scRNA-seq dataset (hMTG-health-sc), both sampled from the healthy human middle temporal gyrus (MTG): and one SRT dataset (hMTG-AD-SRT) coupled with a scRNA-seq dataset (hMTG-AD-sc) from the MTG of Alzheimer’s disease patients (Table 1). This compilation results in a total of fifteen pairs of SRT and scRNA-seq datasets.

LEGEND performs integrated gene clustering on each dataset pair, with resultant clusters assessed in three aspects: 1) the intra-cluster co-expression, proxied by similarity between spatial expression patterns of genes within the same cluster, 2) the co-expression of gene clusters across cell types, and 3) the spatial coherence of gene clusters across tissue domains. Although the true number of gene clusters cannot be determined, clustering performance can still be evaluated since it would show sensible clustering results only if the clustering algorithm works as expected. To control the gene cluster size, we set the target number of clusters to 500 for all tested methods so that on average each cluster has approximately 40 genes, approximating the average number of genes in a typical KEGG pathway [30].

To quantify the similarity of spatial expression patterns between genes within the same cluster, we invent an intra-cluster co-expression quality (ICQ) metric (see Supplementary Text S5). Briefly, we employ the computer vision techniques, Canny Edge Detector and Patch Brightness Detector, to extract and encode edge and brightness traits in a gene’s spatial expression map. The ICQ metric of a cluster is calculated as the average Pearson correlation between the trait encodings of its component genes. We further categorize the gene clusters in each dataset into three groups based on ICQ: high, medium and low. Gene clusters with an ICQ in the top third quantile range are assigned to the high-quality group, those in the bottom third quantile range to the low-quality group, and all remaining clusters to the medium-quality group. Next, a cluster is randomly selected from each of the three groups and we further choose four genes at random from each of the selected clusters. The denoised co-expression patterns of chosen genes are plotted for comparison. As shown in **Figure 2A &2B**, cluster 1, 2, and 3 are selected from high, medium and low co-expression quality groups in the mBrain-SRT dataset and hDLPFC-SRT dataset (slice 151673,). For both datasets, we observe that genes within each cluster exhibit similar spatial expression patterns, particularly for clusters 1 and 2, demonstrating the effectiveness of LEGEND in identifying co-expressed genes.

**Figure 2.**
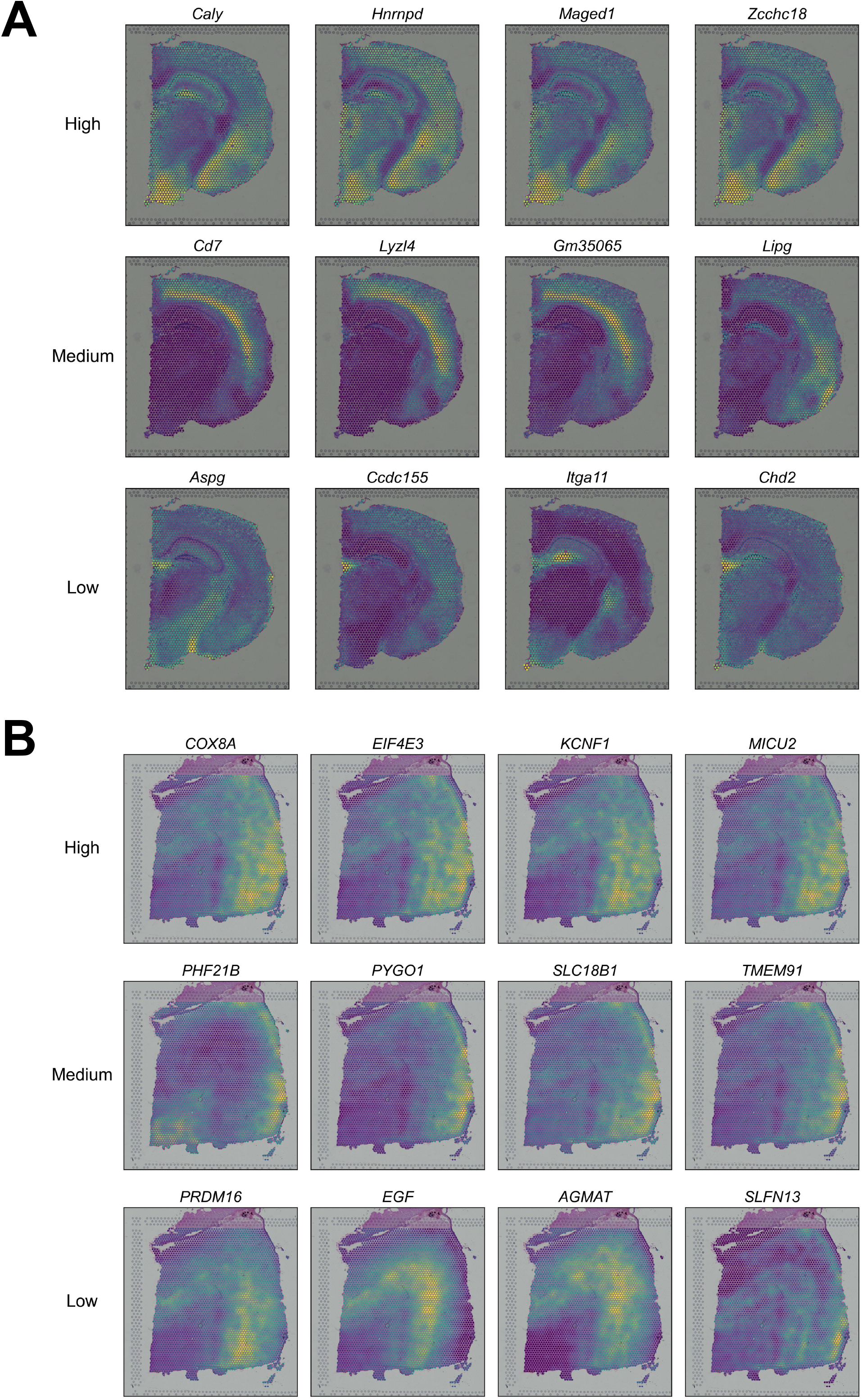
Spatial gene expression patterns of LEGEND-identified gene clusters in mouse brain and human dorsolateral prefrontal cortex (DLPFC). **A**. Mouse brain. Gene clusters identified by LEGEND from the mBrain-SRT dataset are divided into groups of high, medium, and low co-expression quality based on their intra-cluster co-expression quality (ICQ) metrics. Rows represent a gene cluster randomly selected from each of these groups, respectively. The denoised spatial expression patterns of four genes randomly chosen from the cluster are visualized in each row. **B**. Human DLPFC. Analogous to Figure 2A, the denoised spatial patterns of four genes in randomly selected gene clusters identified by LEGEND from the hDLPFC-SRT dataset are visualized.

The DB indices with Pearson distance and weighted Euclidean distance serve as measures for evaluating the overall co-expression and spatial coherence of gene clusters, respectively (see “*Evaluation metrics*” section). Compared to the benchmark methods, LEGEND consistently generates gene clusters with lower values in both types of DB indices (**Figure 3A &3B**) across scRNA-seq and SRT datasets, implying that LEGEND excels in grouping together similar genes while separating dissimilar ones into distinct clusters at both cell type and tissue domain levels, which contributes to a more accurate identification of co-expressed and co-functional gene modules. Moreover, LEGEND’s superior performance over LEGEND-no-sc indicates that integrating sc**/**snRNA-seq with SRT data can effectively enhance gene clustering.

**Figure 3.**
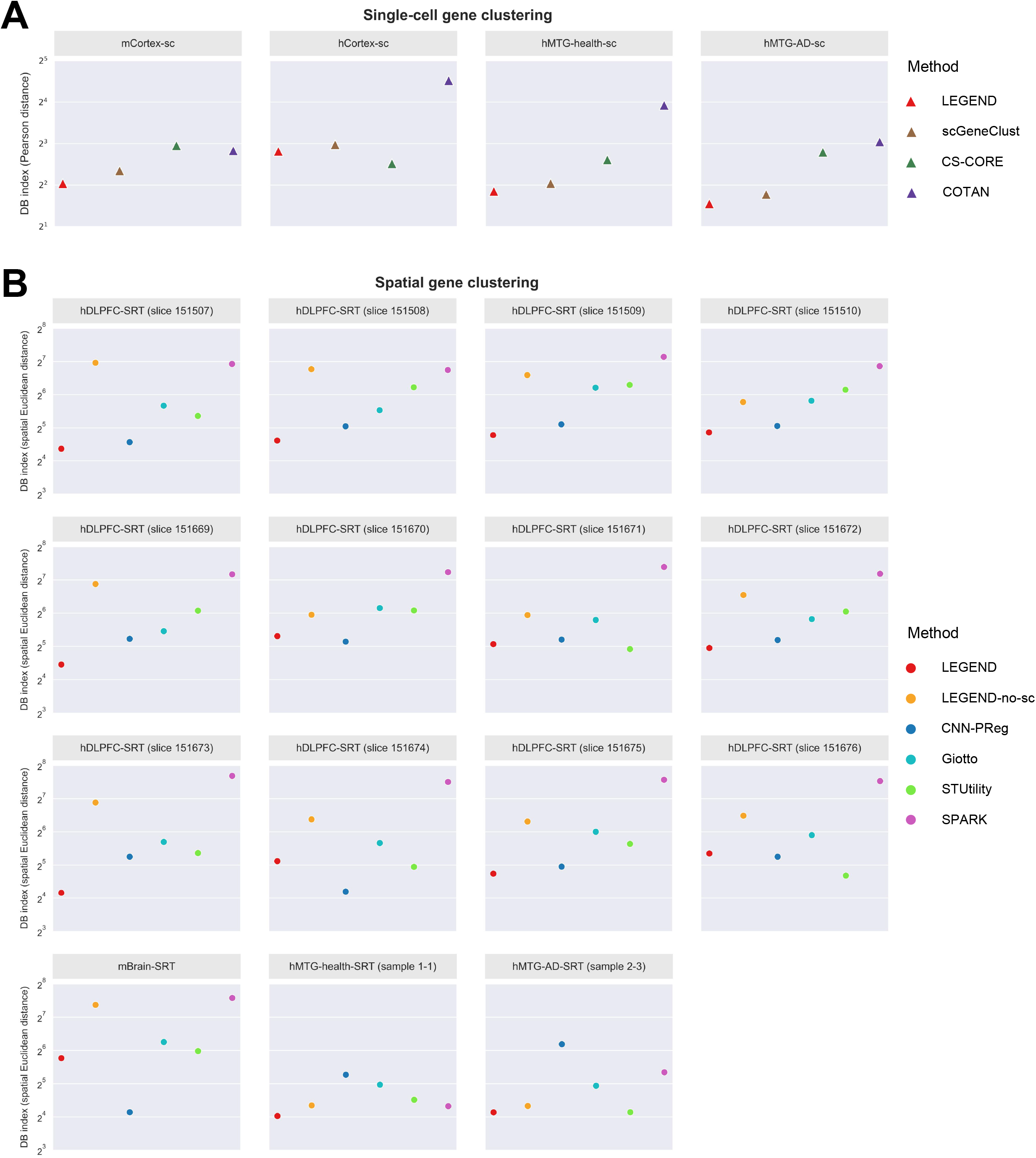
Performance comparison of LEGEND and eight competing gene clustering methods in identifying gene co-expression groups in scRNA-seq and SRT datasets. **A**. The overall qualities of gene co-expression groups identified in each of the four scRNA-seq datasets are quantified using co-expression DB index (y-axis), where a lower score indicates superior clustering quality. **B**. The overall qualities of gene co-expression groups identified in each of the fifteen SRT datasets are quantified using spatial coherence DB index (y-axis), with a lower value indicative of a more effective clustering.

### Biological analyses of LEGEND-generated gene clusters

In this section, we conduct GOBP enrichment, KEGG pathway enrichment and gene co-function analyses on genes within clusters generated by LEGEND to evaluate their biological relevance to the dataset under investigation. This study involves a disease group, which comprises an scRNA-seq (hMTG-AD-sc) and an SRT (hMTG-AD-SRT) dataset of human middle temporal gyrus (MTG) in Alzheimer’s disease (AD), and a normal group comprising an scRNA-seq (hMTG-health-sc) and an SRT (hMTG-health-SRT) dataset of healthy human MTG. The biological analyses are performed on genes from the top relevant gene clusters from both groups (see “*Selection of top relevant gene* section) and a control group with an equivalent number of randomly selected genes. GOBPs and KEGG pathways are defined as AD-related or brain-related based on their connections to the relevant biological process (see Supplementary Text S6).

Here, we only elaborate the results of GOBP enrichment analysis. KEGG pathway enrichment analysis yields similar results (Figure S1 and Table S2). As shown in **Figure 4A** and Table S1, the adjusted p-values of the top 20 most significantly enriched GOBPs of the disease and normal groups are markedly lower than those in the control group, indicating that genes in the disease and normal groups are more likely to participate in the same GOBPs. Additionally, the disease group contains more AD-related GOBPs (15 GOBPs) than the normal (4 GOBPs) and control (2 GOBPs) groups, while the normal group has more brain-related GOBPs (13 GOBPs) than the control group (1 GOBP). These findings demonstrate LEGEND’s capability to identify biologically relevant gene groups the dataset.

**Figure 4.**
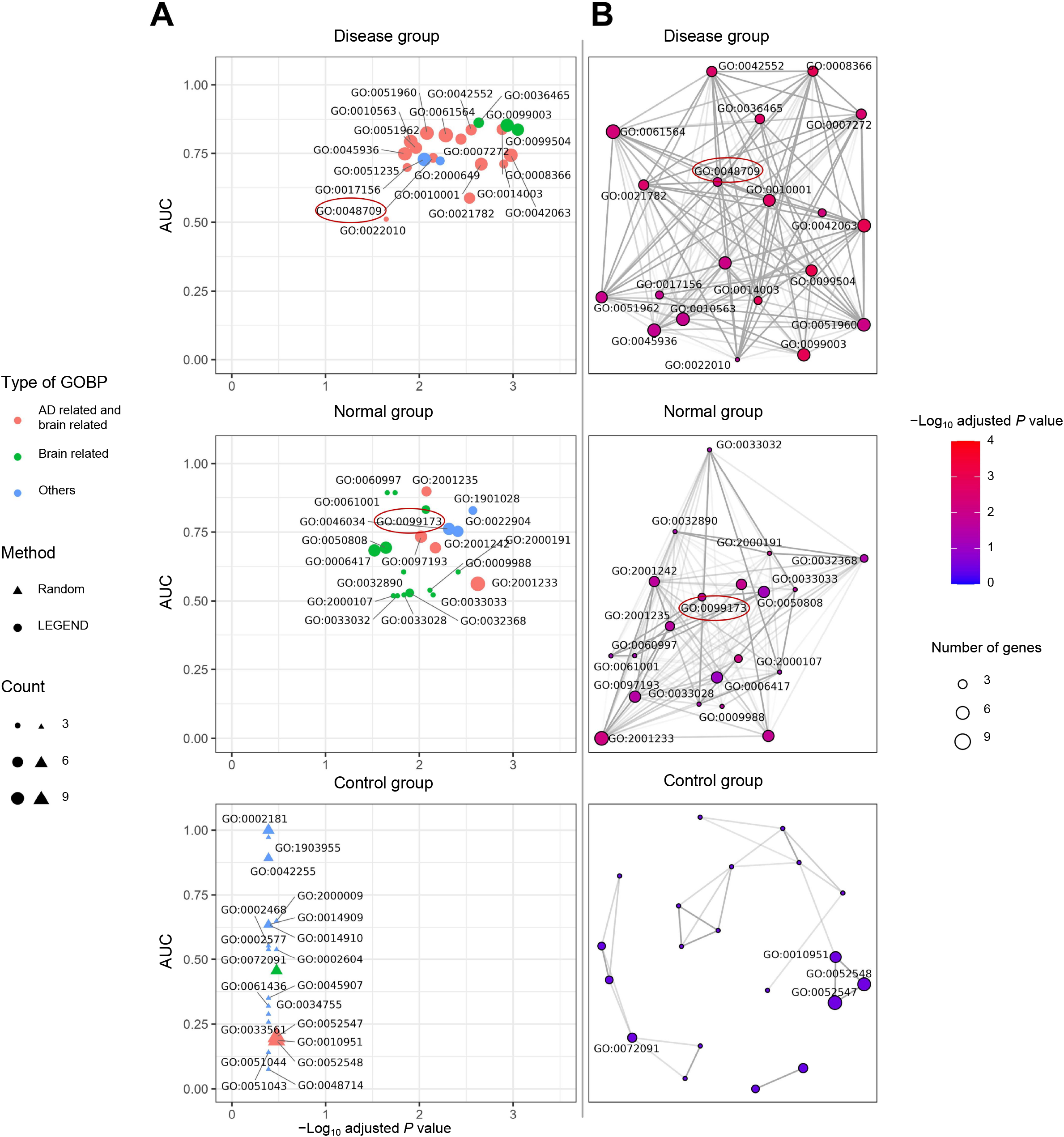
GOBP enrichment and gene cofunction analyses of LEGEND-identified gene clusters. Top relevant gene clusters are collected from datasets of middle temporal gyrus in both healthy and Alzheimer’s disease individuals, forming a normal group and a disease group. A control group comprising a comparable number of randomly selected genes serves as a benchmark. **A**. The 20 most significantly enriched biological processes (BP) for each gene group. The x-axis represents the negative logarithm of adjusted P-values of enrichment significance, while the y-axis represents the AUC scores of the 20 BPs in the gene cofunction analysis, where a higher score indicates stronger cofunctionality within the BP. Red color indicates AD-related (also brain-related) BPs, green color indicates only brain-related BPs, and blue color indicates other BPs. **B**. Enrichment maps of the 20 most significantly enriched pathways for each gene group. The connectivity of the network indicates functional associations among BPs. The node color represents BP’s enrichment significance (negative logarithm of adjusted P-value), with red color indicative of significantly enriched BP. Node size denotes the number of genes involved in the BP.

In the gene co-function analysis, functional coherence among genes in a GOBP is assessed by accuracy of predicting genes’ involvement in that GOBP using their neighboring genes (see “*Gene co-function analysis*” section). Figure 4A reveals that the 20 most significantly enriched GOBPs in both disease and normal groups exhibit notably higher AUC values than those in the control group, supporting that genes in the first two groups tend to be co-functional in the same GOBPs. Figure 4B illustrates that GOBPs enriched in the disease group form a single, densely connected network centering around co-functional elements implicated in the etiology of AD (i.e., “oligodendrocyte differentiation, GO:0048709”), while the GOBPs enriched in the normal group are closely interconnected around co-functional elements related to neural and brain functions (i.e., “postsynapse organization, GO:0099173”). In contrast, GOBPs enriched in the control group form several much smaller, loosely connected networks, most of which lack co-function and are unrelated to AD or brain function. Collectively, these results demonstrate the top relevant clusters generated by LEGEND consist of genes that are co-functional in context-specific GOBPs and pathways.

We delve deeper into the biological significance of these gene clusters by examining the overlapping of the aggregated expression patterns of their component genes with anatomical tissue structure or cell type distributions (see Supplementary Text S7). Six gene clusters identified in the hMTG-AD-SRT dataset exhibit spatial expression patterns closely aligned with the laminar architecture of the human cortex (**Figure 5A**). That is, clusters 1 to 6 overlap with cortical layers I, II, IV, V, VI and WM, respectively. Moreover, their spatial expression patterns overlap with spatial distributions of certain brain resident immune cell types. For instance, higher proportions of astrocytes are found in layer I, neurons (excitatory and inhibitory) in layers II-VI and oligodendrocyte in WM [31], which correspond to the areas where genes of clusters 7, 8 and 9 are over-expressed (Figure 5B). Closer examination reveals that cluster 7 includes marker genes of astrocyte (e.g., *GFAP, ALDH1L1* and *AQP4*), cluster 8 marker genes of neurons (e.g., *GRIN1, GRIN2A* and *GABRA1*), and cluster 9 marker genes of oligodendrocyte (e.g., *OLIG1, CNP* and *PDGFRA*). Figure 5C showcases that the three clusters are respectively enriched for GOBPs relevant to astrocyte (e.g., gliogenesis and glial cell differentiation), neuron (e.g., glutamatergic synaptic transmission and regulation of postsynaptic potential), and oligodendrocyte (e.g., oligodendrocyte differentiation and myelination), further confirming their association with these cell types. More importantly, gene clusters can reveal spatial characteristics of AD pathology—the transcriptional activity of genes in cluster 10, which includes genes related to reactive astrocyte (e.g., *VIM* and *S100B*), significantly increases in WM (Figure 5D), reflecting the spatial distribution of reactive astrocyte in MTG. This observation is consistent with the finding of increased presence of astrocytes in deeper cortical layers and WM in response to the spread of Aβ plagues, neurofibrillary tangles, and neuro-inflammation as AD progresses [32]. Intriguingly, the *TREM2* gene, primarily known for its role in regulating microglial activation and phagocytosis, is also identified in this cluster [33]. As *TREM2* co-expresses with reactive astrocyte-related genes in the same cluster (Figure 5D), it may play a role in regulating reactive astrocytes, a hypothesis also proposed in a recent study [34].

**Figure 5.**
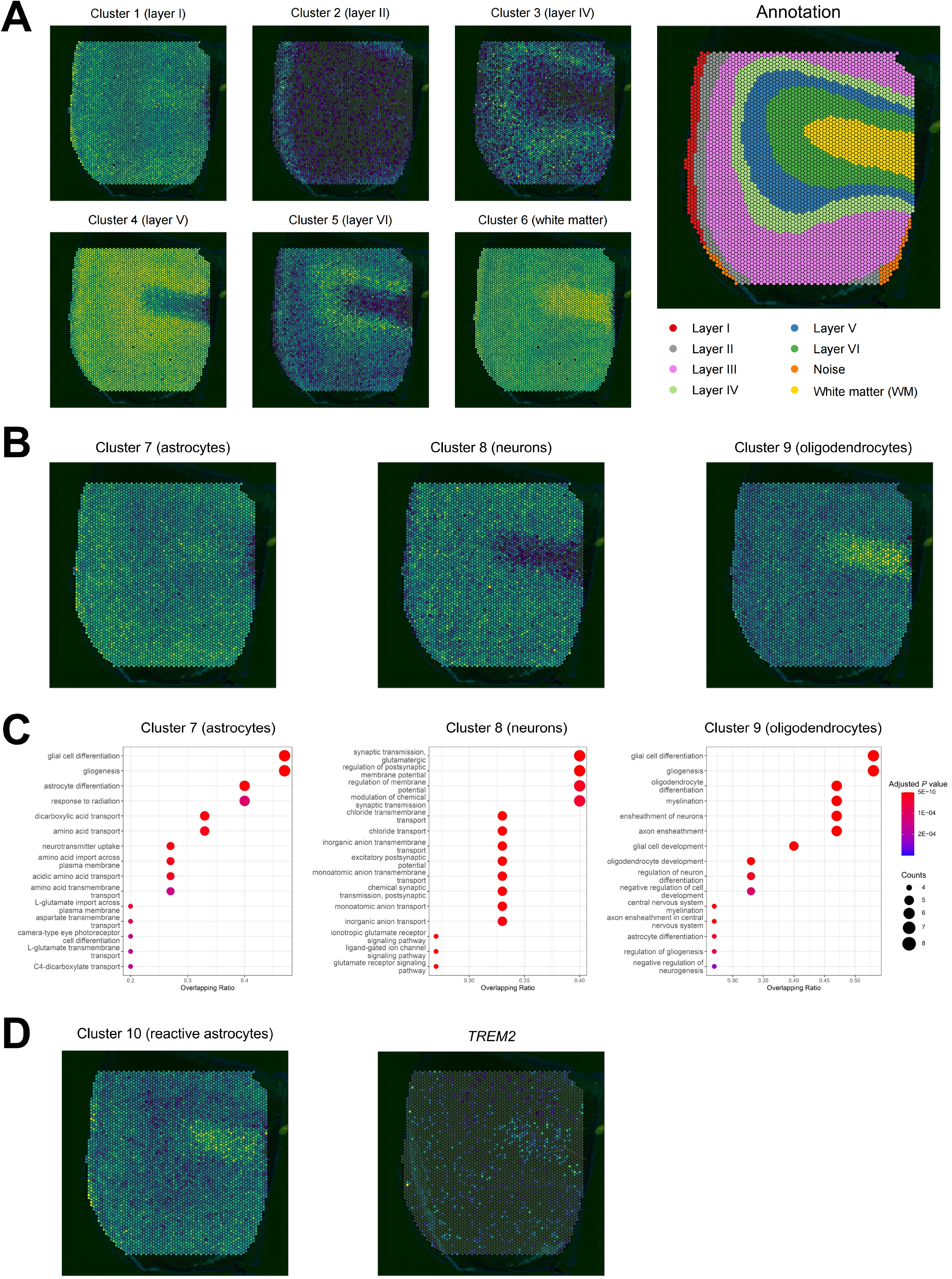
Biological interpretations of LEGEND-identified co-expressed gene clusters in the hMTG-AD-SRT dataset derived from middle temporal gyrus (MTG) of AD patients. **A**. LEGEND-identified gene clusters coincide with the anatomic structure of cortical layers in human brain. The left six panels represent the aggregated spatial expression patterns (module scores) of genes in each of the six gene clusters (clusters 1-6) identified by LEGEND. The right panel displays the annotated anatomical structure of cortical layers. Gene clusters 1-6 overlap with the cortical layer I, II, IV, V, VI and WM, respectively. The brightness in each panel is positively correlated with the aggregated gene expression levels. **B**. LEGEND-identified gene clusters reflect immune cell distributions in cortex layers. The three panels represent the aggregated spatial expression patterns of three LEGEND-identified clusters (clusters 7-9). Genes of clusters 7, 8, and 9 primarily express in cortical layer I, layers II-VI, and WM, respectively. These patterns reflect the typical distributions of astrocytes, neurons, and oligodendrocytes, respectively. **C**. The three panels show the most significantly enriched BPs in gene clusters 7-9. The x-axis represents overlapping ratio, which is the proportion of cluster genes involved in the BP. Circle size indicates the number of genes in the pathway, while red colors indicate significantly enriched BPs. **D**. The left panel showcases a LEGEND-identified gene cluster (cluster 10) whose spatial expression pattern is characterized by significantly elevated transcriptional activity in the WM. This pattern mirrors the pathological distribution of reactive astrocyte in AD brain and is also observed in the spatial expression of the *TREM2* gene (right panel), a cluster 10 member gene and linked to microglial activation in AD.

### Identification of disease-associated gene interactions

In this section, we evaluate LEGEND’s utility in identifying disease-associated gene interactions by exploring shifts in LEGEND-generated gene co-expression networks from healthy state to diseased state. To establish the methodological soundness, we begin by examining the quality of gene interactions implied by the gene partial correlation matrix derived from LEGEND-generated gene redundancy matrix (see “*Identifying disease-associated gene crosstalk*” section). This analysis includes a total number of 7102 genes, forming 25,215,651 gene pairs, among which 8239 are curated transcription factor (TF)-target gene (TG) pairs in the TRRUST v2 database [19,35], serving as the ground truth. LEGEND generates the partial correlation matrix of these genes using a scRNA-seq dataset (hPC-healthy-sc) and a SRT dataset (hDLPFC-SRT, slice 151673), both of which are derived from healthy human prefrontal cortex tissues. Three benchmark methods, including Giotto, CS-CORE, and COTAN, predict gene interactions using the hPC-healthy-sc dataset alone. Each method assigns “scores” to gene pairs to indicate their likelihood of interaction. For instance, in LEGEND, the score is determined by the partial correlation between gene pairs, while in CS-CORE, it is based on a P-value derived from a statistical test under the null hypothesis that two genes are uncorrelated. Gene pairs are then ranked by these scores to select out the most likely interacting gene pairs (i.e., those that are top-ranked) predicted by each method for further analyses. Specifically, CS-CORE selects four gene pair sets of different sizes, containing gene pairs with P-values below thresholds at 1E-02, 5E-03, 1E-03, and 5E-04, respectively. Similarly, each of the other methods also selects four sets of top-ranked gene pairs, with sizes matching the corresponding sets selected by CS-CORE. Across the four sets, LEGEND consistently outperforms the competing methods in recovering the greatest number of TF-TG interactions, as indicated by its superior recall scores shown in **Figure 6A**, which demonstrates LEGEND’s ability in capturing essential gene relationships.

**Figure 6.**
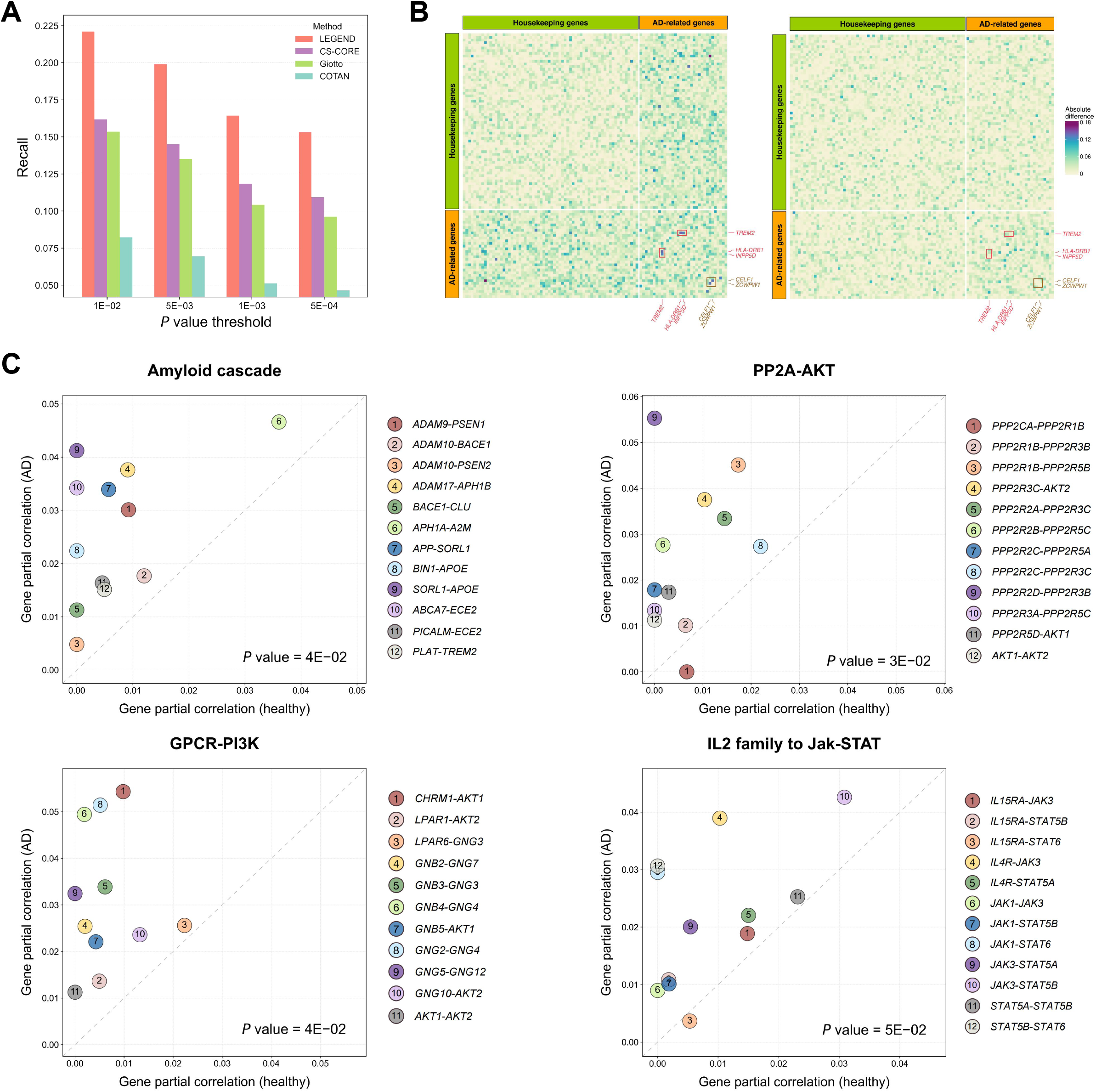
LEGEND reveals disease-associated gene interactions. **A**. Predicting interacting transcriptional factor (TF)-target gene (TG) pairs. Bar groups represent gene pair sets of different sizes (i.e., 23.76%, 21.5%, 17.66%, 16.41%), corresponding to the number of gene pairs with CS-CORE P-values below the specified thresholds (i.e., 1E-02, 5E-03, 1E-03, and 5E-04) on the x-axis. Methods’ effectiveness in recovering known TF-TG interactions is quantified as recall scores displayed on the y-axis. **B**. Heatmaps illustrate the absolute differences in LEGEND-derived partial correlations of 33 AD-associated genes and 66 house-keeping genes between AD versus healthy states (left) and between two healthy conditions (right). Darker shades signify greater shifts. Notably marked with red squares are three gene pairs ({*TREM2, HLA-DRB1*}, {*TREM2, INPP5D*}, {*CELF1, ZCWPW1*}) showing significant correlation shifts in the AD context compared to minor changes between healthy conditions. **C**. Scatterplots depict absolute partial correlations between genes within four established AD-associated gene pathways, including the amyloid cascade, PP2A-AKT, GPCR-PI3K, and Jak-STAT signaling pathways. The vertical and horizontal axes represent partial correlations computed from the AD and healthy states, respectively. Spots in each plot represent gene pairs in the gene signaling pathway. Statistical significance (P-values) of partial correlation discrepancies between the AD state versus the healthy state are noted at the lower-right corner of each plot.

Subsequently, we compile a list of 33 AD-associated genes from previous studies (see Table S3) along with 2178 house-keeping genes cataloged in the HRT Atlas v1.0 database [36], from which 66 genes are randomly selected to form a control gene set. Three partial correlation matrices of these 99 genes, denoted as Φ_*AD*_, Φ_*health*1_, and Φ_*health*2_, are generated by LEGEND from a pair of AD datasets (i.e., hMTG-AD-SRT &hMTG-AD-sc) and two pairs of healthy human MTG datasets (i.e., hMTG-health-SRT &hMTG-health-sc, and hMTG-health-SRT2 &hMTG-health-sc2). Alterations in gene-gene interactions derived from two datasets are measured as absolute value differences between their partial correlation matrices: Φ_*diff1l*_ = |Φ_*AD*_ −Φ_*health*1_| and Φ_*diff*2_ = |Φ _*health*1_ − Φ_*health*2_|.

For easier interpretation, Φ_*diff1*_ and Φ_*diff2*_ are visualized as heatmaps in Figure 6B. For Φ_*diff1*_, we find that gene pairs including at least one AD-associated gene demonstrate significantly greater shifts (Figure 6B, left panel) as opposed to those solely comprising housekeeping genes, implying altered gene interactions involving AD-associated genes in the AD context. In contrast, Φ_*diff2*_ does not show such disparities (Figure 6B, right panel), aligning with expectations that gene relationships among both AD-related and housekeeping genes should remain largely consistent across healthy conditions. For instance, the partial correlations between three gene pairs, including {*TREM2, HLA-DRB1*}, {*TREM2, INPP5D*}, and {*CELF1, ZCWPW1*}, exhibit most significant shifts in the AD context but not between the two healthy states, suggesting their interactions are potentially linked to AD. Intriguingly, recent studies reported that *INPP5D* acts downstream of *TREM2* to regulate the microglial barrier against Aβ toxicity in AD [37]; *TREM2* and *HLA-DRB* families are collectively linked to neuroinflammations throughout the AD continuum [38]; *ZCWPW1* and *CELF1* are co-upregulated in PPARγ knockdown cell-lines associated with AD [39] and in Alzheimer’s Disease risk polymorphisms [40].

We further investigate the altered interactions between genes participating in four established AD-associated gene pathways, including the amyloid cascade [41], PP2A-AKT [42], GPCR-PI3K [42], and Jak-STAT [43] signaling pathways (Table S4), by plotting their partial correlation values in Φ_*AD*_ and Φ_*health*1_ on different axes in scatterplots (Figure 6C). Our analysis reveals that gene pairs within each of the four gene pathways exhibit statistically significant increased correlations in the AD dataset compared to the healthy dataset. These observations underscore LEGEND’s capability in detecting context-specific alterations in gene interactions, thus facilitating the illumination of molecular mechanisms behind diseases.

### Detecting genes with designated spatial expression patterns

Identifying genes with designated spatial expression patterns can be crucial for uncovering genes associated with biological phenotypes and diseases. For example, certain types of brain-resident immune cells (e.g., reactive astrocyte) in AD exhibit specific distribution patterns across cortical layers during the process of neuroinflammation [32]. Exploring genes that display similar patterns in spatial expression provide critical clues to molecular mechanisms driving the spatial distribution of these cells. While numerous methods exist for detecting genes differentially expressed in specific regions [44], they are inadept at identifying genes with varying expression levels across multiple regions. To address this limitation, we introduce an innovative approach capable of simulating a pseudo-gene that mirrors the designated spatial expression patterns (see Supplementary Text S8). By feeding the pseudo-gene alongside the target SRT dataset to LEGEND, real genes resembling the pseudo-gene in spatial expression patterns can be identified via LEGEND-mediated gene clustering (**Figure 7A**).

**Figure 7.**
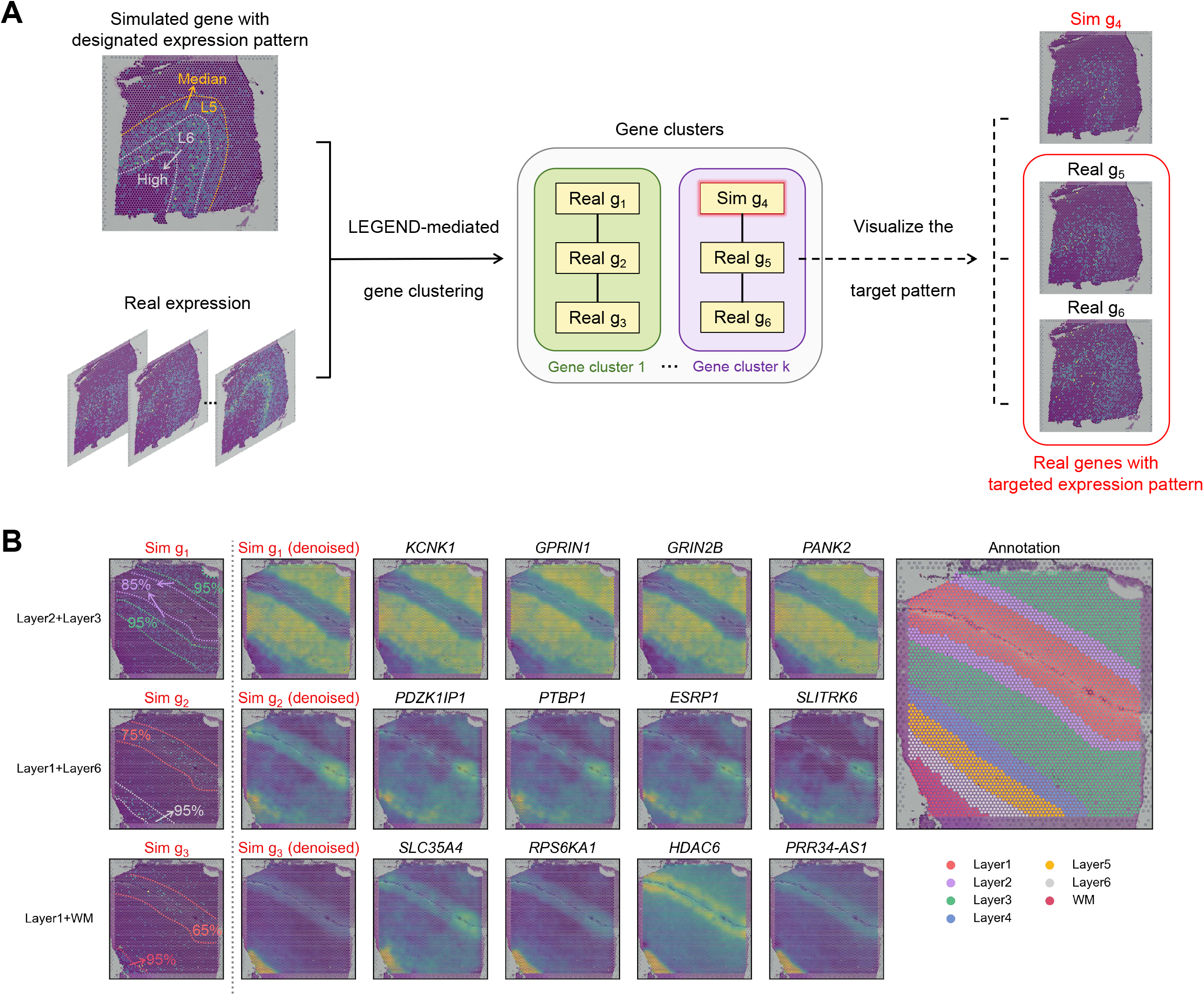
Utilizing LEGEND to pinpoint genes with designated spatial expression patterns. **A**. Workflow overview. This simplified example demonstrates our strategy to identify genes that manifest high expression in cortex layer 6 and median expression in cortex layer 5. A pseudo-gene reflecting these specific expression patterns is simulated and then grouped with real genes through LEGEND-mediated gene clustering. Genes clustered with the pseudo-gene are expected to exhibit the targeted expression patterns. **B**. Application to Human Dorsolateral Prefrontal Cortex: our objective is to identify genes within the hDLPFC-SRT dataset that align with three distinct spatial expression patterns: i) *very-high* expression in layer 3 and *high* in layer 2*i* ii) *very-high* expression in layer 6 and *moderate-high* in layer 1*i* iii) *very-high* expression in WM and *moderate* in layer 1, with all patterns exhibiting *low* expression in other layers. The rightmost panel provides ground-truth cortex layer annotations. The initial column on the left illustrates the simulated pseudo-genes tailored to the desired patterns, followed by a column showing their denoised expressions. The next four columns reveal the denoised expression patterns of four genes from the same cluster as their respective pseudo-genes.

To evaluate this approach, we simulate three pseudo-genes for the hDLPFC-SRT (slice 151509) dataset, each with distinct designated expression patterns across cortex layers of the human DLPFC (Figure 7B). The first pseudo-gene is characterized by a *very-high* expression level in layer 3 and a *high* expression in layer 2: the second exhibits a *very-high* expression level in layer 6 and a *moderate-high* expression in layer 1: the third demonstrates a *very-high* expression level in white matter (WM) and a *moderate* expression in layer 1. Moreover, all three pseudo-genes have a *low* expression level in other layers. Gene expression levels are defined as: “*very-high*” corresponds to the 95th percentile of average expression across all genes, descending through “*high*” at the 85th percentile, “*moderate-high*” at the 75th, “*moderate*” at the 65th, to “*low*” at the 35th percentile. Figure 7B showcases that, from LEGEND-generated gene clusters that contain the simulated pseudo-genes, we successfully pinpoint real genes with spatial expressions matching the corresponding sought-after patterns.

### Improving clustering accuracy of spatial spots and single-cells

LEGEND can improve the information efficiency of feature genes by selecting representative genes with minimized redundancy and maximized relevance from LEGEND-generated gene clusters (see Supplementary Text S9). These genes are used as inputs to downstream algorithms for spatial or single-cell clustering to boost their performances. Here, Leiden and SpaGCN are used as the spatial clustering methods, while Seurat v5 [16] as the single-cell clustering method. Representative genes are selected from gene clusters generated by LEGEND, LEGEND-no-sc or scGeneClust from the 13 pairs of SRT and sc**/**snRNA-seq datasets (Table 1). For comparative purpose, we also include several competing methods for improving feature efficiency, including SpatialDE, BinSpect, and SPARK-X that select spatially variable genes (SVG) in SRT, and Seurat’s VST and M3Drop that select highly variable genes (HVG) in sc**/**snRNA-seq. Each method selects a similar number of genes as input to the clustering algorithms.

We first compare the performance of spatial clustering with LEGEND-selected genes versus with SVGs selected by the competing methods. As shown in **Figure 8A** and Figure S2-S4, LEGEND surpasses the competing methods in enhancing the performance of both SpaGCN and Leiden across the 13 SRT datasets, as indicated by the most improved ARIs and NMIs. Notably, LEGEND’s superiority over scGeneClust and LEGEND-no-sc can be explained as the latter two methods exclude genes only if they are redundant at cell type and tissue domain level, respectively. Conversely, LEGEND only excludes genes redundant at both levels, thereby preserving more complementary information between the two levels. A key observation is that cortex_4 and cortex_5, which are adjacent tissue domains in Figure 8B, can only be accurately differentiated by SpaGCN when using feature genes selected by LEGEND and scGeneClust. We posit that genes capable of discriminating the two domains have relatively invariant expression across tissue domains yet varying expression across cell types. Consequently, they are overlooked by SVG selection methods but not by LEGEND. Similarly, we conduct experiments using the mCortex-sc dataset to compare the performance of single-cell clustering with LEGEND-selected genes versus with HVGs selected by the competing methods. We find that, compared to the competing methods, LEGEND also excels in enhancing Seurat v5’s performance (Figure 8A). These results altogether affirm that LEGEND’s capability in selecting a subset of informational efficient genes that simultaneously capture spatial and cell type variabilities, thus instrumental in both spatial and single-cell clustering.

**Figure 8.**
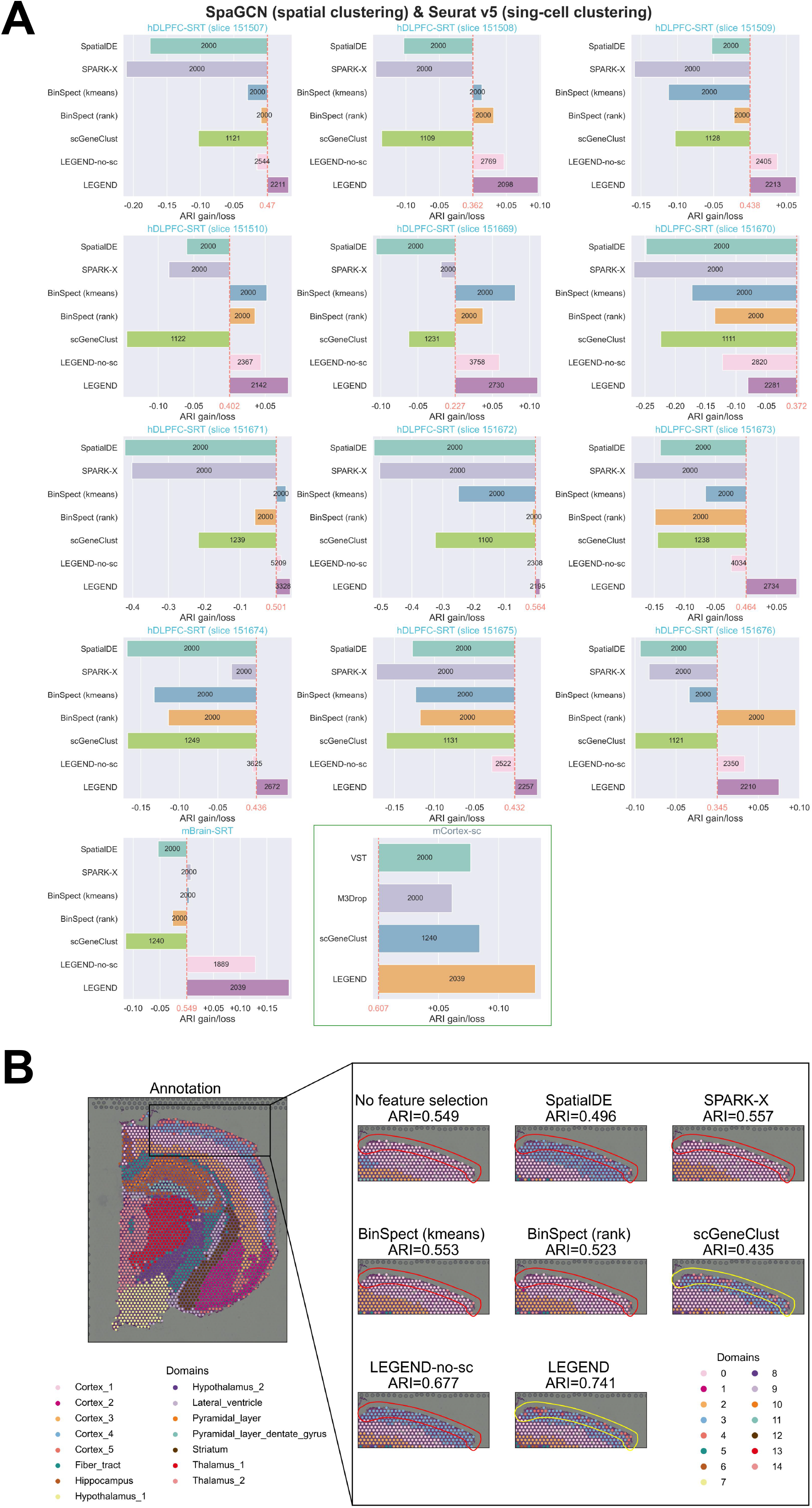
LEGEND improves both single-cell and spatial clustering performance. **A**. SpaGCN is employed for spatial clustering across 13 SRT datasets, while Seurat v5 for single-cell clustering in an scRNA-seq dataset (the rectangle-enclosed panel). Both methods utilize feature gene sets selected by LEGEND or six competing methods. The x-axis displays the ARI changes (+ gain, - loss) compared to baseline performances achieved using the complete gene set (red numbers). The number of genes selected by each method is noted on their bars. **B**. Enhanced tissue domain detection within mouse brain using LEGEND-selected feature gene set. The leftmost panel displays the ground-truth domain labels of the mBrain-SRT dataset, with two adjacent tissue domains, Cortex_4 and Cortex_5, outlined by a black rectangle. The right figures zoom into Cortex_4 and Cortex_5 domains, comparing domain labelling by SpaGCN using feature genes selected by LEGEND and the competing methods. Notably, only scGeneClust and LEGEND effectively distinguish Cortex_5 from Cortex_4, as indicated by yellow circles. The accuracy of spatial domain detection is measured using ARI shown above each right panel.

## DISCUSSION

scRNA-seq and SRT allow the discovery of gene co-expression patterns across cell types and tissue domains, revealing their cofunctionality and interactions in biological systems and diseases. However, existing methods are limited to using either scRNA-seq or SRT data, underutilizing the complementary information between the two data types. This leads to suboptimal cofunctionality within identified gene co-expression groups. To address this limitation, we propose LEGEND, an innovative approach that models gene relationships inherent in paired sc/snRNA-seq and SRT data based on gene relevance, redundancy, and complementarity, based on which a sophisticated gene clustering is employed to identify context-specific gene co-expression groups and gene interactions.

To systematically evaluate LEGEND’s efficacy in identifying biologically meaningful groups of co-expressed genes, we test LEGEND alongside six competing methods across fifteen pairs of SRT and scRNA-seq datasets derived from adult mouse brain and human DLPFC. LEGEND-identified co-expressed gene groups demonstrate not only superior co-expression and spatial coherence over those identified by the competing methods but also significant context-specific biological relevance. This is evidenced by extensive pathway enrichment analyses, gene cofunction analyses, and the congruence of their aggregated spatial expression patterns with known anatomic architecture and cell type distributions. Notably, LEGEND also facilitate the identification of altered gene interactions associated with disease by examining variations in gene interaction networks derived from healthy versus diseased datasets, as suggested by our experiments in which LEGEND successfully reveal altered interactions involving AD-associated genes. Furthermore, we demonstrate LEGEND’s utility in downstream analytical tasks, such as pinpointing genes with designated spatial expression patterns and enhancing the informational efficiency of feature gene set for specific algorithms, e.g., those for spatial and single-cell clustering.

To our knowledge, LEGEND is the first method integrating SRT and sc/snRNA-seq data for identifying groups of co-expressed and cofunctional genes from perspectives of both cell types and spatial tissue domains, alongside a pioneering solution for identifying genes with predefined spatial expression patterns across tissues. LEGEND’s remarkable performance can be attributed to the effective integration of scRNA-seq and SRT data, and a comprehensive quantification of gene informativeness and gene relationships in terms of redundancy, relevance, and complementarity. Nonetheless, there is still rooms for LEGEND’s future improvement. For example, quicker algorithms for calculating gene relevance, redundancy and complementarity could replace the computationally intensive calculations of mutual information involving continuous variables. Lastly, LEGEND-identified gene co-expression groups can potentially serve as genomic “contexts” for the derivation of semantically rich distributed gene representations analogous to the learning of word embeddings from textual contexts in linguistic models.

## Data availability

Our study involves 16 publicly available SRT datasets. The adult mouse brain dataset (mBrain-SRT) is downloaded from the 10x Genomics official website (https://support.10xgenomics.com/spatial-gene-expression/datasets/1.1.0/V1_Adult_Mouse_Brain). Twelve slices of human dorsolateral prefrontal cortex (hDLPFC-SRT) datasets are acquired through spatialLIBD [45] (https://research.libd.org/spatialLIBD/). Three datasets collected from MTG of both AD and healthy individuals (hMTG-AD-SRT sample 2-3, hMTG-health-SRT sample 1-1, and hMTG-health-SRT2 sample 144105) are accessible from the GEO database under accession numbers GSE220442 [12] and GSE226663 [46]. We additionally pair our SRT datasets with six sc/snRNA-seq datasets. The mouse brain cortex dataset (mCortex-sc) is accessible from the GEO database under accession number GSE115746. The human brain cortex dataset (hCortex-sc) is obtained from the Allen Brain Atlas [47] and used in conjunction with hDLPFC-SRT datasets. From the hCortex-sc dataset, we extract cells in MTG to form a new dataset (hMTG-health-sc), which is then paired with the hMTG-health-SRT dataset. Another MTG dataset from a healthy donor (hMTG-health-sc2 sample HC14MTG) is accessible under the GEO accession number GSE188545 [48] and is paired with hMTG-health-SRT2. The hMTG-AD-sc dataset, also from the Allen Brain Atlas, is paired with the hMTG-AD-SRT dataset. The hPC-health-sc (sample NC3) dataset, comprising cells from the prefrontal cortex of a healthy donor, is available under the GEO accession number GSE157827 [49] and is paired with hDLPFC-SRT (slice 151673). LEGEND is freely available as a Python package on Github (https://github.com/ToryDeng/LEGEND).

## Authors’ contributions

XS and HW conceived the idea. XS supervised the study. XS and TD designed the methodology. TD, XS, MH, KX, YX, YL and SC implemented the methods and conducted the computational experiments. XS, TD and YX summarized the results and wrote the manuscript. HW and NX helped revise the manuscript. All authors have read and approved the final manuscript.

## Acknowledgments

The project is funded by Strategic Priority Research Program of Chinese Academy of Sciences (Grant No. XDB38050100) to HW. XS is supported by Excellent Young Scientist Fund of Wuhan City (Grant No. 21129040740).

