## Supplementary figures and images for "LEGEND: Identifying Co-expressed Genes in Multimodal Transcriptomic Sequencing Data"

### Figure S1

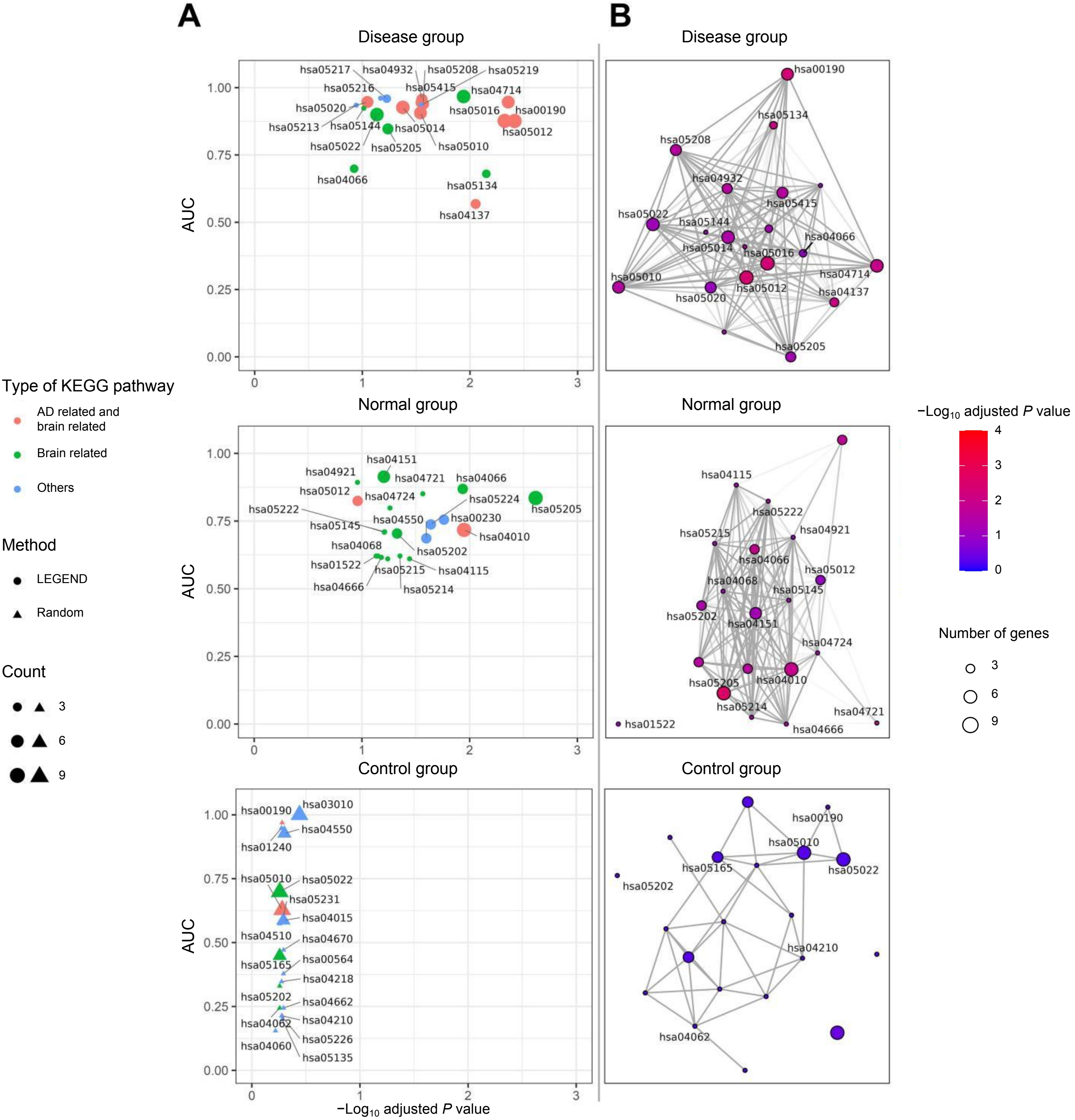

### Figure S2

#### SpaGCN (spatial clustering) & Seurat v5 (sing-cell clustering)

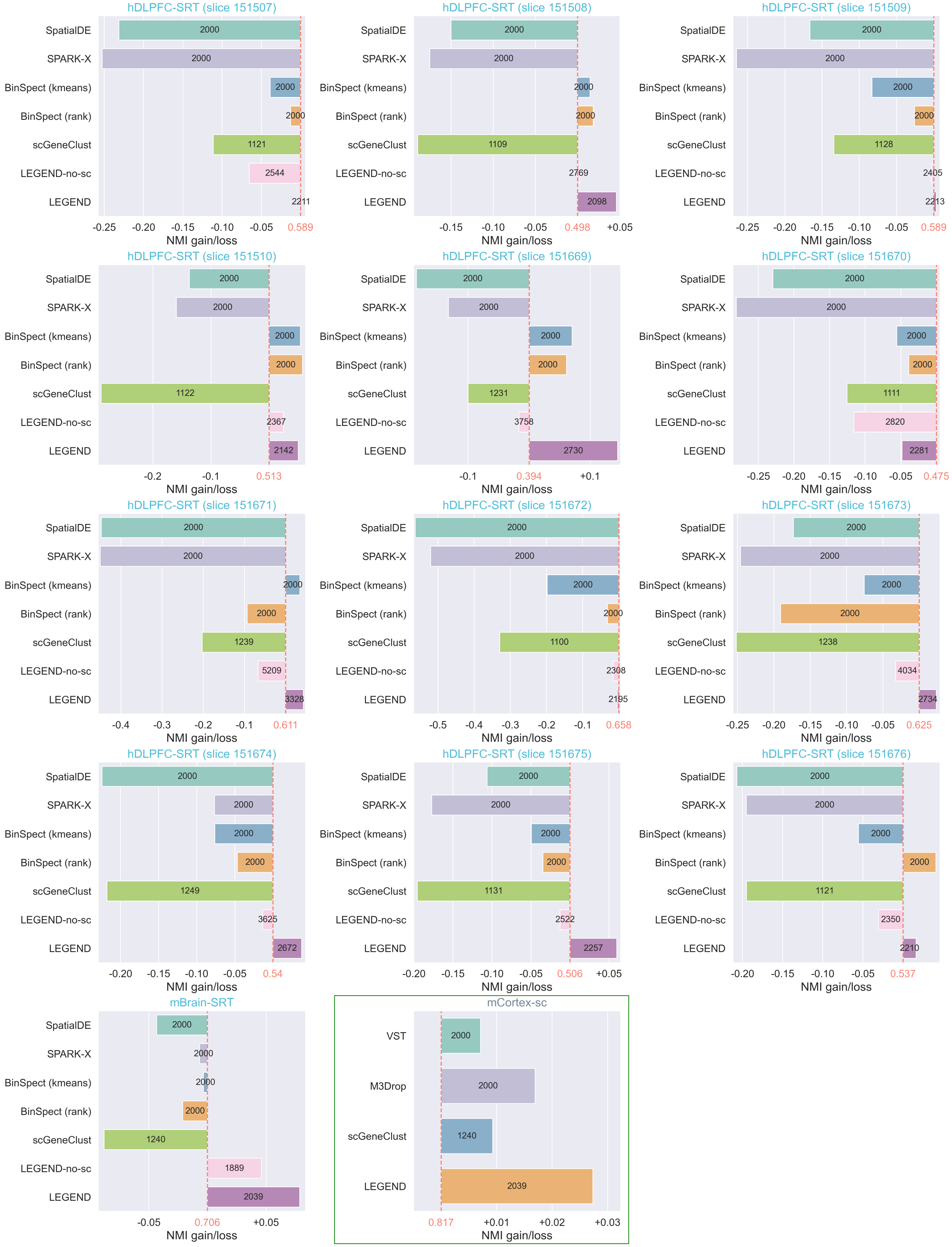

### Figure S3

#### Leiden (spatial clustering)

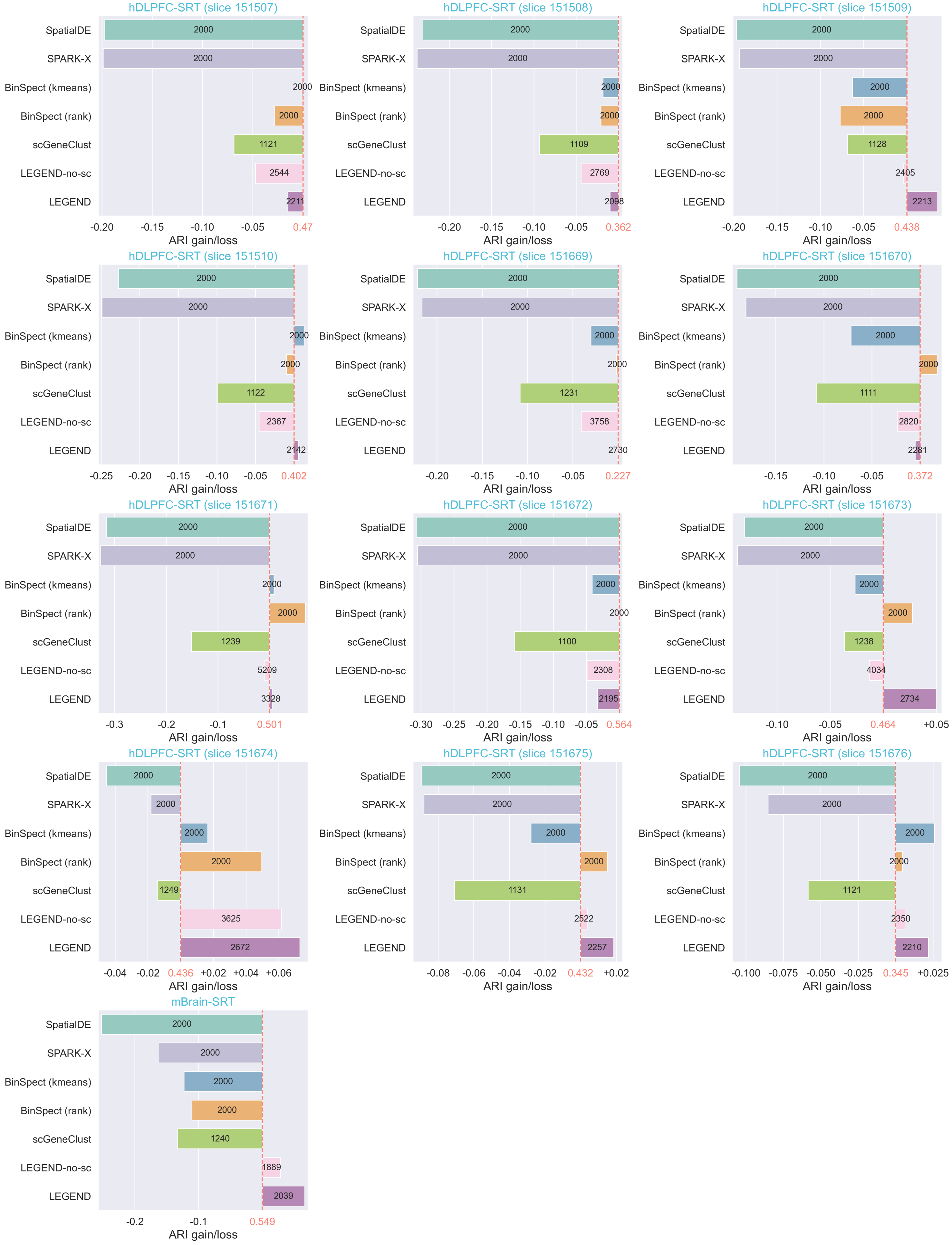

### Figure S4

#### Leiden (spatial clustering)

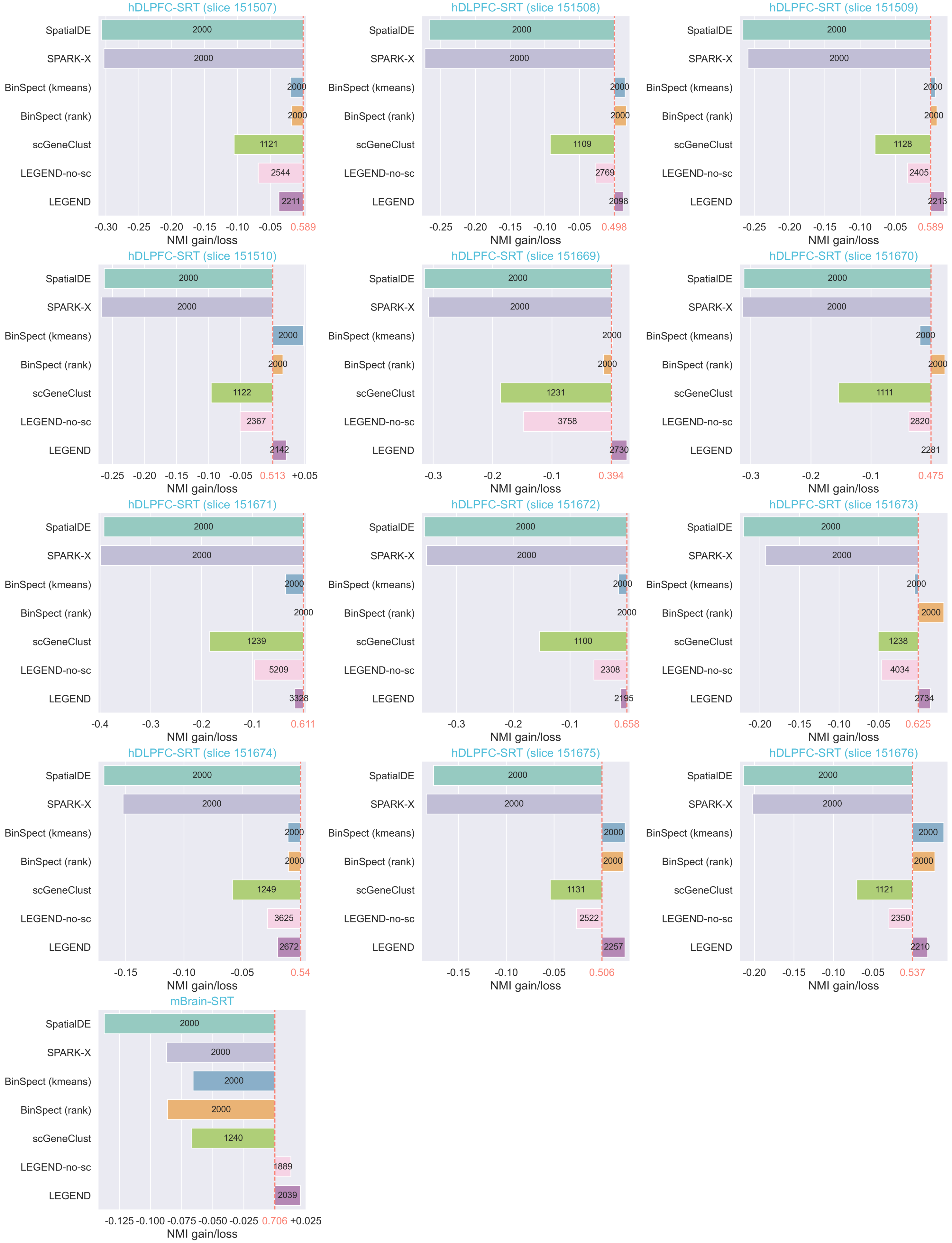
