## Supplementary material for "LEGEND: Identifying Co-expressed Genes in Multimodal Transcriptomic Sequencing Data": Table S1

**Table S1** **List of the 20 most significantly enriched GOBPs in disease, normal and control groups**

| **Group** | **GOBP ID** | **GOBP name** | **Relevance to**  **AD/Brain** |
| --- | --- | --- | --- |
| **Disease**  **Group** | GO:0099003 | vesicle-mediated transport in synapse | Brain Related |
|  | GO:0014003 | oligodendrocyte development | AD and Brain Related |
|  | GO:0010001 | glial cell differentiation | AD and Brain Related |
|  | GO:0036465 | synaptic vesicle recycling | Brain Related |
|  | GO:0042063 | gliogenesis | AD and Brain Related |
|  | GO:0099504 | synaptic vesicle cycle | Brain Related |
|  | GO:0021782 | glial cell development | AD and Brain Related |
|  | GO:0061564 | axon development | AD and Brain Related |
|  | GO:0007272 | ensheathment of neurons | AD and Brain Related |
|  | GO:0010563 | negative regulation of phosphorus metabolic process | AD and Brain Related |
|  | GO:0008366 | axon ensheathment | AD and Brain Related |
|  | GO:0045936 | negative regulation of phosphate metabolic process | AD and Brain Related |
|  | GO:0017156 | calcium-ion regulated exocytosis | AD and Brain Related |
|  | GO:0042552 | myelination | AD and Brain Related |
|  | GO:2000649 | regulation of sodium ion transmembrane transporter activity | Others |
|  | GO:0048709 | oligodendrocyte differentiation | AD and Brain Related |
|  | GO:0051235 | maintenance of location | Others |
|  | GO:0051962 | positive regulation of nervous system development | AD and Brain Related |
|  | GO:0051960 | regulation of nervous system development | AD and Brain Related |
|  | GO:0022010 | central nervous system myelination | AD and Brain Related |
| **Normal**  **Group** | GO:1901028 | regulation of mitochondrial outer membrane permeabilization involved in apoptotic signaling pathway | Others |
|  | GO:2001233 | regulation of apoptotic signaling pathway | AD and Brain Related |
|  | GO:0033033 | negative regulation of myeloid cell apoptotic process | Brain Related |
|  | GO:2001242 | regulation of intrinsic apoptotic signaling pathway | AD and Brain Related |
|  | GO:0097193 | intrinsic apoptotic signaling pathway | AD and Brain Related |
|  | GO:0022904 | respiratory electron transport chain | Others |
|  | GO:0046034 | ATP metabolic process | Others |
|  | GO:2001235 | positive regulation of apoptotic signaling pathway | AD and Brain Related |
|  | GO:2000191 | regulation of fatty acid transport | Brain Related |
|  | GO:0061001 | regulation of dendritic spine morphogenesis | Brain Related |
|  | GO:0032890 | regulation of organic acid transport | Brain Related |
|  | GO:0033032 | regulation of myeloid cell apoptotic process | Brain Related |
|  | GO:0099173 | postsynapse organization | Brain Related |
|  | GO:0060997 | dendritic spine morphogenesis | Brain Related |
|  | GO:2000107 | negative regulation of leukocyte apoptotic process | Brain Related |
|  | GO:0050808 | synapse organization | Brain Related |
|  | GO:0033028 | myeloid cell apoptotic process | Brain Related |
|  | GO:0006417 | regulation of translation | Brain Related |
|  | GO:0009988 | cell-cell recognition | Brain Related |
|  | GO:0032368 | regulation of lipid transport | Brain Related |
| **Control**  **Group** | GO:0052547 | regulation of peptidase activity | AD and Brain Related |
|  | GO:0052548 | regulation of endopeptidase activity | AD and Brain Related |
|  | GO:0010951 | negative regulation of endopeptidase activity | AD and Brain Related |
|  | GO:0072091 | regulation of stem cell proliferation | Brain Related |
|  | GO:0002604 | regulation of dendritic cell antigen processing and presentation | Others |
|  | GO:2000009 | negative regulation of protein localization to cell surface | Others |
|  | GO:0002468 | dendritic cell antigen processing and presentation | Others |
|  | GO:0051044 | positive regulation of membrane protein ectodomain proteolysis | Others |
|  | GO:0042255 | ribosome assembly | Others |
|  | GO:0002577 | regulation of antigen processing and presentation | Others |
|  | GO:0034755 | iron ion transmembrane transport | Others |
|  | GO:0051043 | regulation of membrane protein ectodomain proteolysis | Others |
|  | GO:0048714 | positive regulation of oligodendrocyte differentiation | Others |
|  | GO:0002181 | cytoplasmic translation | Others |
|  | GO:0061436 | establishment of skin barrier | Others |
|  | GO:0014910 | regulation of smooth muscle cell migration | Others |
|  | GO:0033561 | regulation of water loss via skin | Others |
|  | GO:1903955 | positive regulation of protein targeting to mitochondrion | Others |
|  | GO:0014909 | smooth muscle cell migration | Others |
|  | GO:0045907 | positive regulation of vasoconstriction | Others |
