## Supplementary material for "LEGEND: Identifying Co-expressed Genes in Multimodal Transcriptomic Sequencing Data": Table S2

**Table S2 List of the 20 most significant enriched KEGG pathways in disease, normal and control groups**

| **Group** | **KEGG**  **pathway ID** | **KEGG**  **pathway name** | **Relevance to**  **AD/Brain** |
| --- | --- | --- | --- |
| **Disease**  **Group** | hsa00190 | Oxidative phosphorylation | AD and Brain Related |
|  | hsa05012 | Parkinson disease | AD and Brain Related |
|  | hsa04137 | Mitophagy - animal | AD and Brain Related |
|  | hsa04714 | Thermogenesis | Brain Related |
|  | hsa05208 | Chemical carcinogenesis - reactive oxygen species | AD and Brain Related |
|  | hsa05016 | Huntington disease | AD and Brain Related |
|  | hsa05014 | Amyotrophic lateral sclerosis | AD and Brain Related |
|  | hsa05216 | Thyroid cancer | Others |
|  | hsa04932 | Non-alcoholic fatty liver disease | AD and Brain Related |
|  | hsa05217 | Basal cell carcinoma | Others |
|  | hsa05219 | Bladder cancer | Others |
|  | hsa05415 | Diabetic cardiomyopathy | AD and Brain Related |
|  | hsa05134 | Legionellosis | Brain Related |
|  | hsa05213 | Endometrial cancer | Others |
|  | hsa05144 | Malaria | Brain Related |
|  | hsa05205 | Proteoglycans in cancer | Brain Related |
|  | hsa05020 | Prion disease | AD and Brain Related |
|  | hsa04066 | HIF-1 signaling pathway | Brain Related |
|  | hsa05022 | Pathways of neurodegeneration - multiple diseases | Brain Related |
|  | hsa05010 | Alzheimer disease | AD and Brain Related |
| **Normal**  **Group** | hsa05205 | Proteoglycans in cancer | Brain Related |
|  | hsa04066 | HIF-1 signaling pathway | Brain Related |
|  | hsa04010 | MAPK signaling pathway | AD and Brain Related |
|  | hsa00230 | Purine metabolism | Others |
|  | hsa05224 | Breast cancer | Others |
|  | hsa04550 | Signaling pathways regulating pluripotency of stem cells | Others |
|  | hsa04721 | Synaptic vesicle cycle | Brain Related |
|  | hsa04115 | p53 signaling pathway | Brain Related |
|  | hsa05214 | Glioma | Brain Related |
|  | hsa05202 | Transcriptional misregulation in cancer | Brain Related |
|  | hsa04666 | Fc gamma R-mediated phagocytosis | Brain Related |
|  | hsa05215 | Prostate cancer | Others |
|  | hsa05145 | Toxoplasmosis | Brain Related |
|  | hsa01522 | Endocrine resistance | Brain Related |
|  | hsa04151 | PI3K-Akt signaling pathway | Brain Related |
|  | hsa05222 | Small cell lung cancer | Brain Related |
|  | hsa05012 | Parkinson disease | AD and Brain Related |
|  | hsa04724 | Glutamatergic synapse | Brain Related |
|  | hsa04068 | FoxO signaling pathway | Brain Related |
|  | hsa04921 | Oxytocin signaling pathway | Brain Related |
| **Control**  **Group** | hsa03010 | Ribosome | Others |
|  | hsa04550 | Signaling pathways regulating pluripotency of stem cells | Others |
|  | hsa04662 | B cell receptor signaling pathway | Others |
|  | hsa05231 | Choline metabolism in cancer | Others |
|  | hsa00564 | Glycerophospholipid metabolism | Others |
|  | hsa04015 | Rap1 signaling pathway | Others |
|  | hsa04670 | Leukocyte transendothelial migration | Others |
|  | hsa05010 | Alzheimer disease | AD and Brain Related |
|  | hsa00190 | Oxidative phosphorylation | AD and Brain Related |
|  | hsa04210 | Apoptosis | AD and Brain Related |
|  | hsa05135 | Yersinia infection | Others |
|  | hsa05226 | Gastric cancer | Others |
|  | hsa01240 | Biosynthesis of cofactors | Others |
|  | hsa04218 | Cellular senescence | Others |
|  | hsa05022 | Pathways of neurodegeneration - multiple diseases | Brain Related |
|  | hsa05165 | Human papillomavirus infection | Brain Related |
|  | hsa04062 | Chemokine signaling pathway | Brain Related |
|  | hsa05202 | Transcriptional misregulation in cancer | Others |
|  | hsa04510 | Focal adhesion | Others |
|  | hsa04060 | Cytokine-cytokine receptor interaction | Others |
