## Supplementary material for "LEGEND: Identifying Co-expressed Genes in Multimodal Transcriptomic Sequencing Data": Table S3

**Table S3 List of AD-associated genes**

| **AD-associated gene(s)** | **Reference** |
| --- | --- |
| APP | [1] |
| PSEN1, PSEN2 | [2] |
| TREM2 | [3] |
| SORL1 | [4] |
| ABCA7 | [5] |
| ATXN1 | [6] |
| CHRNA4 | [7] |
| MAPT | [8] |
| CST3 | [9] |
| TF | [10] |
| GAB2 | [11] |
| APOE, BCR, CTSS | [12] |
| PLD3, UNC5C, ADAM10, CLU, BIN1, CD2AP, PICALM, HLA-DRB1, INPP5D, MEF2C, PTK2B, ZCWPW1, CELF1, FERMT2, SLC24A4, RIN3 | [13] |
| IQCK, WWOX | [14] |
