## Supplementary material for "LEGEND: Identifying Co-expressed Genes in Multimodal Transcriptomic Sequencing Data": Table S4

**Table S4 List of AD-associated gene pathways**

| **Pathway** | **Genes involved in the pathway** | **Reference** |
| --- | --- | --- |
| the amyloid cascade signaling pathway | ADAM9, ADAM10, ADAM17, BACE1, PSEN1, PSEN2, APH1A, APH1B, APP, BIN1, SORL1, APOE, ABCA7, CLU, PICALM, A2M, ECE2, PLAT, TREM2 | [1] |
| PP2A-AKT signaling pathway | PPP2CA, PPP2R1B, PPP2R2A, PPP2R3C, PPP2R2B, PPP2R2C, PPP2R2D, PPP2R3A, PPP2R3B, PPP2R5A, PPP2R5B, PPP2R5C, PPP2R5D, AKT1, AKT2 | [2] |
| GPCR-PI3K signaling pathway | CHRM1, LPAR1, LPAR6, GNB2, GNB3, GNB4, GNB5, GNG2, GNG3, GNG4, GNG5, GNG7, GNG10, GNG12, AKT1, AKT2 | [2] |
| IL2 family to Jak-STAT signaling pathway | IL15RA, IL4R, JAK1, JAK3, STAT5A, STAT5B, STAT6 | [3] |
