## Supplementary Text for "LEGEND: Identifying Co-expressed Genes in Multimodal Transcriptomic Sequencing Data"

### Text S1. Pseudo labelling of cells in unannotated sc/snRNA-seq datasets

Given the accessibility of annotated scRNA-seq dataset, we assume single-cell labels are available by default. In case of unavailable annotated scRNA-seq data, we propose the following approach that groups together cells that are highly likely to be in the same clusters, the indices of which are used as highly confident pseudo-labels. The algorithm includes the following steps:

Step 1: GMM-based consensus clustering. Ten Gaussian mixture models (GMMs) are trained using the first 2 to 11 gene-level principal components (PCs) of the expression matrix to group cells into clusters, each generating the posterior probabilities that cells belong to their respective clusters. The number of gene-level PCs used to train GMMs can be determined by the elbow point on the curve of cumulative explained variance ratio. The number of clusters $\nu$ for the GMM can be determined by existing methods [1–3]. Let $\mathbf{L}^{k}\in{\mathbb{\mathbb{N}}}^{n}$ and $\mathbf{P}^{k}\in{\mathbb{\mathbb{R}}}^{n}$ denote the cluster indices and posterior probabilities of all $n$ cells in the $k^{\text{th}}$ GMM clustering. A binary $n\times n$ co-membership matrix $\mathbf{M}^{k}$ in the $k^{\text{th}}$ GMM clustering can be defined as:

$$\begin{matrix} \mathbf{M}_{ij}^{k}=\left\{ \begin{matrix} 1, & \text{if}\mathbf{L}_{i}^{k}=\mathbf{L}_{j}^{k}\wedge min\left( \mathbf{P}_{i}^{k},\mathbf{P}_{j}^{k} \right)\geq0.95 \\ 0, & \text{otherwise} \end{matrix} \right. \end{matrix}$$

That is, if we are confident enough that cells $i$ and $j$ belong to the same cluster in the $k^{\text{th}}$ GMM clustering, $\mathbf{M}_{ij}^{k}$ is given a value of 1, otherwise 0. All co-membership matrices generated from the 10 GMM clustering are summed up to form a single matrix $\mathbf{W}=\sum_{k=1}^{10} \mathbf{M}^{k}$, in which $\mathbf{W}_{ij}$ represents the frequency that cells $i$ and $j$ are confidently believed to be of the same type. We then define a threshold $\alpha$ (default value 8) to set the values to 0 for the cell pairs that are not consistently in the same group across all GMMs:

$$\begin{matrix} \mathbf{W}_{ij}=\left\{ \begin{matrix} \mathbf{W}_{ij}, & \text{if}\mathbf{W}_{ij}\geq\alpha\\ 0, & \text{otherwise} \end{matrix} \right. \end{matrix}$$

Step 2: Clustering and pseudo-labelling of HC cells. $\mathbf{W}$ in the above step is used to construct a weighted undirected graph, where graph nodes represent cells and $\mathbf{W}_{ij}$ represents the edge weight between node $i$ and $j$. Leiden algorithm is applied on the graph to cluster cells connected by heavy-weighted edges. The top-$m$ largest-sized clusters consisting of cells that belong to the same type with high confidence are preserved as HC clusters. $m$ is determined as:

$$m=min\left( \nu,\left| \{\tilde{C}\left| \tilde{C}\in\tilde{C}_{leiden}, \right|\tilde{C}|\geq n_{1}\} \right| \right)$$

Here $\nu$ represents the estimated number of clusters in GMM clustering, $\tilde{C}_{leiden}$ represents the set of cell clusters generated by Leiden. $\left| \cdot\right|$ denotes the cardinality of a set and $n_{1}\in{\mathbb{\mathbb{N}}}^{+}$ is a predefined number (default 10).

### Text S2. Calculation of mutual information involving continuous variables

In our study, we leverage a k-nearest neighbor based algorithm to estimate the mutual information between two continuous variables (e.g., redundancy) or between a continuous variable and a discrete variable (e.g., relevance) [4,5]. Let $\mathrm{MI}\left( X,Y \right)$ denote the mutual information between two variables $X$ and $Y$, $H\left( \cdot\right)$ denote the entropy of a variable, we have:

$\begin{matrix} MI\left( X,Y \right) & =H\left( X \right)+H\left( Y \right)-H\left( X,Y \right) (1a) \\ & =H\left( Y \right)-H\left( Y|X \right) (1b) \end{matrix}$

Briefly, the algorithm first leverages the Kozachenko-Leonenko estimator [6] to estimate a probability density $\mu_{i}$ for each observation $x_{i}$. For $x_{i}$, it calculates the distance from its k-th nearest neighbor, denoted by $\delta_{i}$, then calculates the probability of observing exact $K$ neighbors within $\delta_{i}$ via a trinomial formula to obtain $\mu_{i}$. Then, the entropy $H\left( x_{i} \right)$ within $x_{i}$’s k-nearest neighbor space can be calculated as the expected value of $\ln\mu_{i}$ which is a function of $K$, $N$ (number of observations) and $\delta_{i}$, and $H\left( X \right)\approx\left[ \sum_{i=1}^{N} H\left( x_{i} \right) \right]/N$. $H\left( Y \right)$, $H\left( X,Y \right)$, $H\left( Y|X \right)$ can be approximated in a similar way. $\mathrm{MI}\left( X,Y \right)$ is calculated according to equation 1a in case of two continuous variables, or equation 1b in case of one continuous and one discrete variable.

### Text S3. Proof of Theorem 1

*Proof.* When $\mathbf{Red}\left( g_{i},g_{j} \right)<\mathbf{Comp}\left( g_{i},g_{j} \right)$, we have the following transformation and inequality:

$$\begin{matrix} \mathbf{Red}\left( g_{i},g_{j} \right)-\mathbf{Comp}\left( g_{i},g_{j} \right)= & \mathrm{MI}\left( g_{i},g_{j} \right)-MI\left( g_{i},g_{j}|\boldsymbol{L} \right) \\ = & H\left( g_{i} \right)-H\left( g_{i}|g_{j} \right)-H\left( g_{i}|\boldsymbol{L} \right)+H\left( g_{i}|g_{j},\boldsymbol{L} \right) \\ = & \mathrm{MI}\left( g_{i}|\boldsymbol{L} \right)-MI\left( g_{i},\boldsymbol{L}|g_{j} \right) \\ < & 0 \end{matrix}$$

### Text S4. Fine-tuning of gene clusters with EOM algorithm

In the MST, two genes connected by an edge with a low RRI value are not sufficiently redundant relative to their complementarity/relevance, making them less likely to be co-functional. Consequently, their edge should be severed to separate them into different gene clusters. The EOM algorithm is leveraged to convert the MST into a hierarchy of gene clusters, represented as a dendrogram, by partitioning the MST at different RRI threshold levels. Specifically, the RRI threshold starts with a low value at the top of the hierarchy, so all genes in the MST belong to a single cluster, which is the root node of the dendrogram. As the RRI threshold increases in a top-down manner, fewer MST edges possess large enough RRI values to be preserved, leading to an increasing number of smaller gene clusters toward the hierarchy’s bottom. In cases where cutting an edge results in two child clusters—one small and the other of a comparable size to their parent cluster— the small child cluster is considered as a noise cluster and thus should be merged back into its parent cluster. We set a minimum cluster size of five for a cluster not to be considered as a noise one. After constructing the cluster tree, MST splits are examined in a top-down manner. An edge split is considered to truly generate new clusters only when both new clusters have at least five genes. This process simplifies the cluster tree, which then only consists of persistent gene clusters as branches that gradually lose their genes as the RRI increases.

To further select “high quality” gene clusters from the hierarchy, we define the stability coefficient for a cluster $i$ within the hierarchy as:

$$SC\left( i \right)=\sum_{p\in cluster i} \left( \mathrm{RRI}_{p}-\mathrm{RRI}_{\mathrm{birth}} \right)$$

Here, $\mathrm{RRI}_{\mathrm{birth}}$ and $\mathrm{RRI}_{p}$ correspond to the RRI threshold level when cluster $i$ splits off from its parent cluster and becomes its own cluster, and the RRI threshold level when gene p falls out of cluster $i$. The EOM algorithm proceeds as follows: Starting with the leaf nodes as selected clusters, the algorithm traverses up the cluster tree. For a cluster $i$, if the sum of its children’s stability coefficients is greater than its own, then $SC\left( i \right)$ is updated to be the sum value. Otherwise, cluster $i$ is considered as a sufficiently stable cluster and thus selected, while all its descendant clusters are unselected. The algorithm continues until it reaches the root node, outputting the set of selected clusters as the final result.

### Text S5. Intra-cluster co-expression quality (ICQ) metric

To evaluate the spatial expression similarity between genes within the same cluster, we propose an intra-cluster co-expression quality (ICQ) metric based on computer vision techniques. Specifically, we apply the Canny Edge Detector (also known as Canny operator) and Patch Brightness Detector to the normalized spatial expression matrix $\mathbf{X}^{\boldsymbol{u}}\boldsymbol{\in}{\mathbb{\mathbb{R}}}^{N_{x}\times N_{y}}$ of gene $u$, where $N_{x} and N_{y}$ represent number of spatial spots along the horizontal and vertical directions of the spatial map. Out-of-tissue spatial spots are all padded with 0s in$\mathbf{X}^{\boldsymbol{u}}$.

We first smoothen $\mathbf{X}^{\boldsymbol{u}}$ with a convolutional Gaussian kernel, obtaining a denoised expression matrix ${\tilde{\boldsymbol{X}}}^{\boldsymbol{u}}$. The Canny Edge Detector is then applied on ${\tilde{\boldsymbol{X}}}^{\boldsymbol{u}}$ to generate an edge trait matrix $\mathcal{C}^{\boldsymbol{u}}$, which reflects the expression continuity of gene $u$. We apply the Patch Brightness Detector by segmenting ${\tilde{\boldsymbol{X}}}^{\boldsymbol{u}}$into small patches containing $\boldsymbol{k}\boldsymbol{\times}\boldsymbol{k spots}, where$ $\boldsymbol{k}\in\{1,3,5\}$, from whichthree patch brightness mean matrices $\mathcal{A}^{\boldsymbol{u}}\left( \boldsymbol{k} \right)$and two variance matrices $\mathcal{S}^{\boldsymbol{u}}\left( \boldsymbol{k} \right)\boldsymbol{(}eligible only for \boldsymbol{k=}3,5\boldsymbol{)}$ are calculated:

$$\mathcal{A}_{i,j}^{u}\left( k \right)= \mathrm{avg}\left( X_{s\left( k \right),t\left( k \right)}^{u} \right),$$

$$\mathcal{S}_{i,j}^{u}\left( k \right)= \mathrm{std}\left( X_{s\left( k \right),t\left( k \right)}^{u} \right),$$

$$1\leq i\leq N_{x},1\leq j\leq N_{y}$$

$$s(k)=\left[ i-\frac{\left( k-1 \right)}{2}:i+\frac{\left( k-1 \right)}{2} \right],t(k)=\left[ j-\frac{\left( k-1 \right)}{2}:j+\frac{\left( k-1 \right)}{2} \right]$$

Finally, the ICQ matrix $S^{\mathcal{c}}$ of cluster $\mathcal{c}$is calculated as the average Pearson correlation between its gene pairs’ $\mathcal{C}^{\boldsymbol{u}}$, $\mathcal{A}^{\boldsymbol{u}}\left( \boldsymbol{k} \right)$ and $\mathcal{S}^{\boldsymbol{u}}\left( \boldsymbol{k} \right)$:

$S_{u,v}^{\mathcal{c}}=\left\{ \begin{aligned} \mathrm{avg}\left( \rho_{u,v}\left( \Xi^{u},\Xi^{v} \right) \right);\Xi\in\{\mathcal{A}\left( k \right)\mathcal{,S}\left( k \right)\mathcal{,C}\}, k\in1,3,5,u\neq v \\ 0, u=v \end{aligned} \right.$and $S^{\mathcal{c}}\in{\mathbb{\mathbb{R}}}^{d\times d}$

Here, $u,v$ represent any two genes within cluster $\mathcal{c}$, while $d$ denotes the number of genes within the cluster $\mathcal{c}$.

**Text S6.** **Defining AD-related and brain-related GOBPs and KEGG pathways**

Child terms of “generation of neurons (GO:0048699)”, “neuron death (GO:0070997)”, “neurogenesis (GO:0022008)”, “apoptotic process (GO:0006915)” and “axonogenesis (GO:0007409)” in the GO database are defined as AD-related, given their close relationship to AD pathology. GOBPs associated with other neural activities, such as “dendritic spine morphogenesis (GO:0060997)” and their respective child terms, are categorized as brain-related, reflecting general neural-related processes. For KEGG pathways, those that are reported to share cellular or molecular mechanisms with the AD (e.g., “hsa00190: Oxidative phosphorylation”) are categorized as AD-related, while pathways that have connections with other brain functions are categorized (e.g., “hsa04921: Oxytocin signaling pathway”) as brain-related.

**Text S7.** **Visualization of clusters’ aggregated gene expression patterns**

To visualize the aggregated gene expression patterns of a cluster on the SRT spatial map, we employ the “AddModuleScore” function from the Seurat R package to calculate module scores across all spatial spots for that cluster. Specifically, all genes of the cluster are binned based on their average expression, and an equal number of control genes are randomly selected from each bin. The differences between the average expressions of the set of cluster genes and the set of control genes serve as the cluster’s module scores across all spatial spots. Finally, the clusters’ module scores at every spatial spot are colored and displayed on the SRT spatial map, using a color spectrum that corresponds to the range of the module scores. This approach allows us to better understand how the spatial distribution of gene expression relates to the anatomical tissue structure.

**Text S8.** **Simulate designated gene spatial expression patterns**

We introduce a generalized linear model (GLM)-based method to simulate a pseudo-gene with desired spatial expression patterns, based on which real genes with similar expression patterns can be identified via LEGEND-mediated gene clustering. This task is not trivial—on one hand, given no prior knowledge about which real genes exhibit the target expression pattern, existing reference-based simulation methods, as seen in Andersson et al^38^., fail to pinpoint an appropriate gene for reference. On the other hand, without referring to real target data, reference-free simulation methods like SRTsim^39^ fall short of generating ST data with consistent data properties.

Our method allows specifying precise gene expression levels in targeted tissue regions. Specifically, let $G$ represent the set of all genes and $S$ the set of all spots in the target dataset. The size-factor-normalized read counts of any gene $i\in G$ at spot $j \in S$, $X_{i,j}$, follows a negative binomial distribution:

$X_{i,j}\sim NB\left( \mu_{i},s_{i} \right)$,

where $\mu_{i}$ and $s_{i}$ are mean and dispersion parameters, respectively. Then we have the following equations:

$$\sigma_{i}^{2}=\mu_{i}+\frac{\mu_{i}^{2}}{s_{i}}\Longrightarrow\text{cv}_{i}^{2}=\frac{1}{\mu_{i}}+\frac{1}{s_{i}}, (2)$$

where $\sigma_{i}^{2}$ and $\text{cv}_{i}^{2}$ represent variance and squared coefficient of variation (SCV) of gene $i$, respectively. As proved in Brennecke et al^40^., we have:

$$E\left[ {\hat{\text{cv}}}_{i}^{2} \right]\approx a_{0}+\frac{a_{1}}{\hat{\mu}_{i}},$$

$$\hat{\mu}_{i}=\frac{\sum_{j} X_{i,j}}{N},$$

$${\hat{\text{cv}}}_{i}^{2}=\frac{\sum_{j} \left( X_{i,j}-\hat{\mu}_{i} \right)^{2}/\left( N-1 \right)}{\hat{\mu}_{i}^{2}},$$

where ${\hat{\text{cv}}}_{i}^{2}$ and $\hat{\mu}_{i}$ are sample SCV and mean, respectively. $N$ denotes the total number of spots in the ST dataset. Note that ${\hat{\text{cv}}}_{i}^{2}$ approximately follows a $\chi^{2}$ or gamma distribution. By taking $\frac{1}{\hat{\mu}_{i}}$ as input, we can use a GLM of the gamma family with an identity link function to perform the regression:

$$\sum_{i} \log\left( P\left( {\hat{\text{cv}}}_{i}^{2} | \alpha,\beta_{i} \right) \right)=\sum_{i} \left( -\alpha log\left( \beta_{i} \right)-\frac{{\hat{\text{cv}}}_{i}^{2}}{\beta_{i}}+\left( \alpha-1 \right)\log\left( {\hat{\text{cv}}}_{i}^{2} \right) \right)+C_{1},$$

$$E\left[ {\hat{\text{cv}}}_{i}^{2} | \alpha,\beta_{i} \right]=\alpha\beta_{i}=\eta_{i}=a_{0}+\frac{a_{1}}{\hat{\mu}_{i}},$$

where $\alpha=\frac{1}{\sigma}$ is a constant, and $\sigma$ denotes the GLM dispersion parameter. Let the natural parameter of the GLM be $\theta_{i}=\frac{-1}{E\left[ {\hat{\text{cv}}}_{i}^{2} | \alpha,\beta_{i} \right]},$ then we have:

$$\sum_{i} \log\left( P\left( {\hat{\text{cv}}}_{i}^{2} | \alpha,\beta_{i} \right) \right)\propto\sum_{i} \left( \frac{\theta_{i}{\hat{\text{cv}}}_{i}^{2}-log\left( -\frac{1}{\theta_{i}} \right)}{\sigma}+\frac{1-\sigma}{\sigma}\log\left( {\hat{\text{cv}}}_{i}^{2} \right) \right),$$

from which coefficients $a_{0}$ and $a_{1}$ can be estimated using the Fisher-scoring method.

Assume we aim to simulate a gene with a top $t\%$expression level in a spatial region $S_{1}\in S$, and the gene conforms to a NB distribution with mean $\mu$ and dispersion $s$. We first calculate and rank ascendingly the average expressions of all genes in the target dataset, denoted as $L$. Then, $\mu$ and the corresponding squared coefficient of variation, $\text{cv}$, can be estimated as:

$$\mu=L@t\%,$$

$$\text{cv}^{2}\approx E\left[ \text{cv}^{2} \right]=\alpha_{0}+\frac{\alpha_{1}}{\mu}.$$

According to equation 2, $s$ can be computed as:

$$s=\frac{1}{\text{cv}^{2}-\frac{1}{\mu}}.$$

Let $N_{1}$ denote the number of spots within $S_{1}$ region. For each spot $s_{j}\in S_{1}$, the median gene expression across all genes is calculated, based on which $S_{1}$ is reordered ascendingly as $\tilde{S}_{1}$. Next, we simulate $N_{1}$ observations from $NB\left( \mu,s \right)$, which are sorted in ascending order and assigned to correspondingly ranked spots in $\tilde{S}_{1}$. This ordered assignment helps to preserve the typical spatial gene expression structure inherent in the ST dataset. Thereby, we obtain a simulated gene expressed at the top $t\%$ quantile level within the $S_{1}$ region. We can repeat this procedure to make the simulated gene to be expressed at specified quantile levels within arbitrary tissue regions.

**Text S9.** **Representative gene selection**

As the genes within the same gene cluster have redundant information, it is beneficial to select a few representative genes from them and discard the rest to reduce feature redundancy and computational load in downstream analytical tasks such as spatial clustering of spatial spots. In our case, we choose genes that rank within the top 20% in their respective clusters based on the $\mathbf{Re}\mathbf{l}^{\mathrm{com}}$ metric as representative genes, which represent the most discriminative subset of genes for tissue domains or cell types within each cluster.
